# Novel Epidermal Oxysterols Function as Alarm Substances in Zebrafish

**DOI:** 10.1101/2023.09.26.559639

**Authors:** Yaxi Li, Zhi Yan, Ainuo Lin, Xiao Yang, Xiaodong Li, Xiuli Yin, Weiming Li, Ke Li

## Abstract

Aquatic animals often use chemical cues to signal predation risk. When injured, shoaling fish skins release alarm substances that induce intense fear and a suite of anti-predator behaviors in other shoal members. However, the chemical nature of alarm substances remains largely unknown. Here we show that zebrafish alarm substance comprises 24-methyl-5α-cholestane-3α,7α,12α,24,28-pentahydroxy 28-sulfate, a novel oxysterol sulfate, and 5α-cyprinol sulfate. These compounds are present in zebrafish skin extract and, at concentrations of less than one nanomolar, each induces anti-predator behaviors and increases cortisol levels. Their mixture, at its natural ratio, replicates the skin extract in eliciting the full suite of anti-predator behavior patterns. Our findings reveal a molecular-level mechanism whereby fish signal predation danger.

## Main

Many animals use alarm cues for defense against predators. These cues may be visual, auditory, or chemical. Chemical alarm cues are common among social insects and aquatic animals. These chemicals, also known as alarm substances or alarm pheromones, are signal molecules essential to survival^1,2^. Alarm substances warn conspecifics and sometimes other members of the group to flee, shoal, seek refuge, or fortify defenses^2^. Most known alarm pheromones have been isolated from terrestrial social insects such as ants, bees, and aphids. These identified pheromones are volatile compounds that dissipate efficiently in the air^3,4^. The existence of alarm substances in fish was first proposed in 1938 by von Frisch, who inferred the presence of *schreckstoff* (also known as “fear substance”) from behavioral responses of minnows to injured conspecifics^5^. Subsequently, extensive studies have shown that these substances elicit stereotypical behaviors and physiological responses in Ostariophysan^6^, a superorder that includes over 10,000 fish species. In addition, several fish species are found to release disturbance cues when threatened (but not injured)^7,8^. There is also evidence that fish detect odorants of potential predators and consequently become disturbed^8,9^. It appears that fish, the largest group of vertebrate animals, rely heavily on chemical cues from various sources to assess predation risk.

For decades, a great number of researchers have sought to identify the chemicals that cause “fear” or disturbance in fish^10^. In black tetra and giant danio, hypoxanthine-*N*-oxide (H3NO) was postulated as the alarm substance at excessively high concentrations^11^; however, it has not been detected reliably in the skin^12^. Mathuru et al. (2012)^13^ noticed that vigorous shaking causes zebrafish to release substances that induce a mild fright reaction. They further showed that, through fractionation of skin extract and behavioral assays, glycosaminoglycan chondroitins, major components of mucus, induce some components of alarm behavior, although the activation level is lower than those induced by skin extracts^13^. Their evidence indicated that alarm cues and possibly disturbance cues are present in zebrafish skin tissues and that the full complement of alarm substances has yet to be characterized.

We sought to identify alarm substance components that induce the same anti-predator responses induced by skin extract in zebrafish. When exposed to skin extracts, zebrafish show distinct and conspicuous locomotion patterns, including high-velocity erratic swimming, bottom-dwelling, and freezing^14^. Protocols for measuring these behaviors have been well established^15^. Low concentrations of conspecific blood (0.01%) also elicit robust defensive responses in adult zebrafish, indicating that damage-released chemical cues in the blood and epidermis mediate anti-predator behaviors^16^. In addition, cortisol levels are elevated in zebrafish exposed to chemical cues from injured or disturbed individuals of the same species^17^. Based on the original and revised definitions of alarm pheromone^2^, and the characteristics of alarm substance examined in numerous fish species^1,8,9^, we reasoned that the zebrafish alarm substance should be composed of a chemical or a mixture of chemicals that is found in skin tissues, induces a full complement of anti-predator behaviors and a stress response, and shows expected functions at concentrations consistent with the abundance found in skin extract^18^. We further speculated that the active components may be substances that diffuse rapidly in water, mirroring the known insect alarm pheromones that often dissipate rapidly in the air.

We used a fractionation strategy guided by behavioral assays^15^ and tracked the alarm substance activities of skin extract to two molecules. These two candidate components of alarm substance were isolated by chromatography and characterized unequivocally by nuclear magnetic resonance spectroscopy (NMR) and ultra-high performance liquid chromatography–high-resolution mass spectrometry (UPLC–HRMS). When mixed at the ratio found in the skin extract and diluted to concentrations expected from the skin extract, the two compounds induced the full suite of anti-predator behaviors and increased cortisol levels in adult zebrafish.

## Results

### Skin extract (SKE) induced anti-predator behaviors

To develop a reproducible bioassay that measures the full suite of anti-predator behaviors, we validated the known behavioral responses elicited by skin extract in zebrafish using a novel tank diving paradigm, a well-accepted protocol (Supplementary Fig. 1)^15,17^. Naïve zebrafish 3-5 months old were each used once in the assay. Skin extract (SKE) was prepared from skin samples of ∼800 zebrafish. Test stimulants were prepared by dissolving 5% of SKE in water to make a stock solution equivalent to the skins of 80 fish/mL, which was serially diluted by 10–10^6^ fold (annotated as 10×SKE–10^6^×SKE) and tested for their behavioral activities (Supplementary Fig. 1–2)^14,19-21^. Twelve potential anti-predatory behavior parameters and derived indices of zebrafish were recorded and analyzed (Supplementary Fig. 3). The results showed that four parameters characteristic of predator avoidance behaviors in zebrafish^14,22^, namely erratic movement duration (%), time in bottom half (%), freezing duration (%), and latency to upper half (s) (Supplementary Fig. 3a-d), increased in fish exposed to the test stimulants. These four parameters were therefore selected to track the anti-predator activities of SKE fractions and to confirm the function of purified components. Recorded examples of erratic movement and freezing behaviors were given in video 1&2. Notably, the 10^3^-fold dilution of SKE (1000×SKE; equivalent to skin extract from 0.08 fish/L in the test tank) induced each of the four selected behaviors (Supplementary Fig. 3a–d), consistent with previous findings^14^. Thus, we used 1000×SKE as the benchmark (Fig. 1b–e) to track the active components in bioassay-guided experiments.

**Fig. 1.**
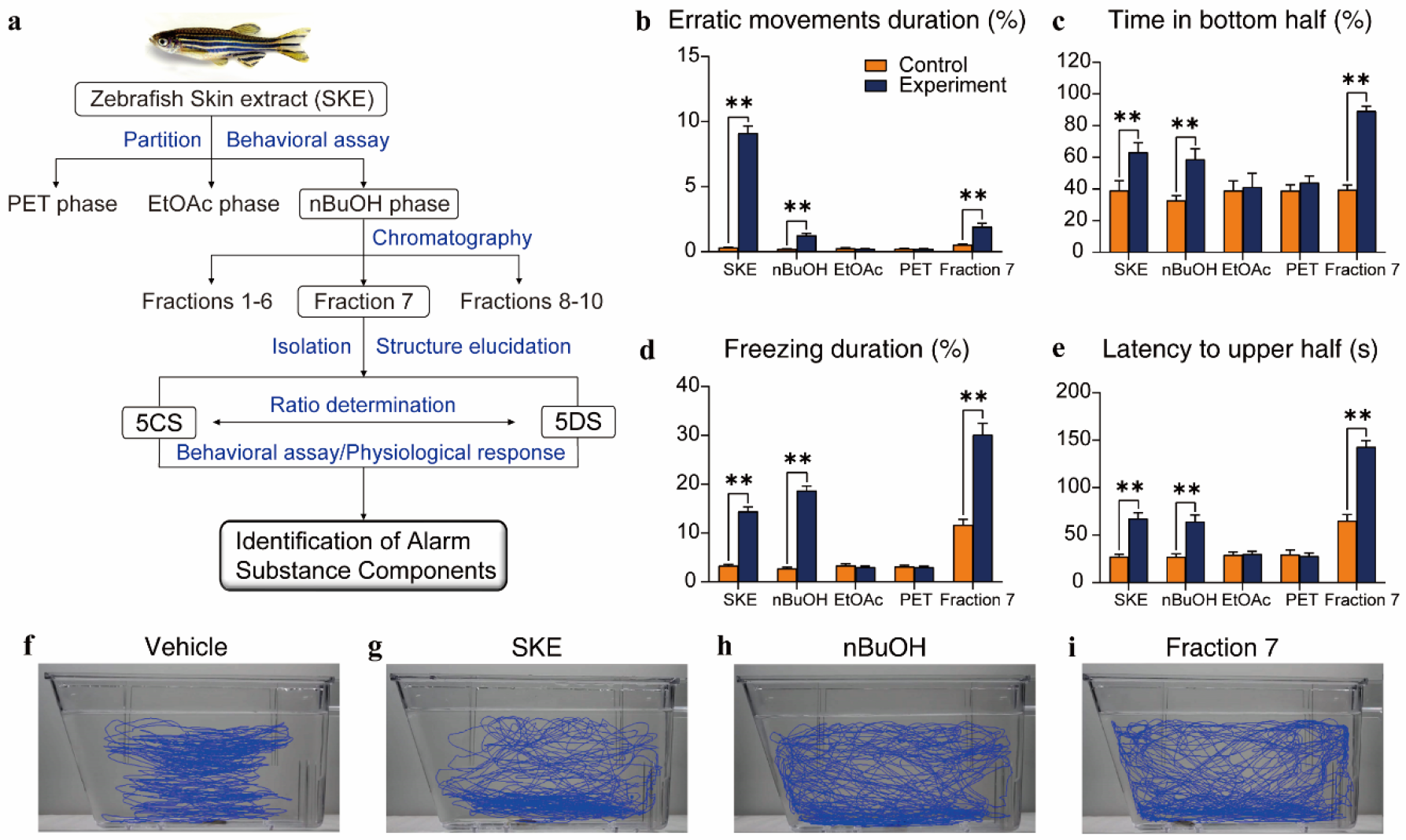
**a)** Bioassay-guided fractionation to identify alarm substance components. Boxed fractions replicated the activity of skin extract (SKE) in inducing changes in four behavioral phenotypes, including **b**) erratic movement duration (%), **c**) time in the bottom half (%), **d**) freezing duration (%), and **e**) latency to the upper half (s). Fractions of successive separation by petroleum (PET), ethyl acetate (EtOAc), n-butanol (nBuOH), and fraction 7 of nBuOH, were reconstituted in water to equivalents of a 1000-fold dilution of SKE. Representative motion traces of females exposed to **f**) vehicle, **g**) SKE, **h**) nBuOH, **i)** fraction 7, respectively. *, *P* < 0.05; **, *P* < 0.01. See Supplementary Fig. 2 for detailed fractionation procedures. In **b**-**e**, the exact sizes of biologically independent samples and the results of independent sample t-tests or Mann-Whitney U tests are summarized in Supplementary Table 1.

### The polar fraction of SKE contained the alarm substance components

To fractionate alarm substance components (Fig. 1a), SKE (80% of the sample) was successively extracted with petroleum ether (PET), ethyl acetate (EtOAc), and n-butanol (nBuOH) to segregate compounds of increasing polarity. Each extract was reconstituted in water at a concentration equivalent to 1000×SKE and assayed for its behavioral activities (Figs. 1b–d & Supplementary Fig. 2). When presented to adult zebrafish, the nBuOH extract replicated SKE in inducing erratic movement duration (%) (Fig. 1b), time in bottom half (%) (Fig. 1c), freezing duration (%) (Fig. 1d), and latency to upper half (s) (Fig. 1e), whereas neither EtOAc nor PET extract induced these behaviors (Fig. 1b–e). These results confirmed that the active components are relatively hydrophilic.

We fractionated the nBuOH extract by semi-preparative HPLC with a reverse-phase column, on which the major components coeluted in fraction 7. Fraction 7 induced the same four behaviors (Fig. 1b–e), replicating the behavioral activities of 1000×SKE and the nBuOH extract.

The behavioral traces (Fig. 1f–i) also showed SKE, nBuOH, and fraction 7 induced changes in the movement trajectories. Behavioral trajectories stimulated by SKE (Fig. 1g) showed an apparent bottom-dwelling (time in bottom half) response as compared to those induced by the vehicle (Fig. 1f), a pattern that was also observed in fish exposed to nBuOH and fraction 7 (Fig. 1h–i). Compared to those induced by the vehicle, the traces induced by SKE, nBuOH, and fraction 7 suggest the test fish touched the tank side walls with increasing frequencies (Figs. 1h–i). This may be because the alarm substance components were gradually purified as the remaining interfering substances were eliminated, causing the zebrafish to respond more strongly to the stimuli. Touching the side walls likely reflects evasive behaviors upon exposure to the treatment stimulatns. Taken together, fraction 7 contained the alarm substance candidates.

### Alarm substance candidates were identified as two oxysterols

We identified a predominant peak consisting of two compounds within fraction 7 (Fig. 2a and insets). Using HPLC-HRMS, we further isolated and detected them. The NMR spectra and HRMS spectra indicated the compound at a mass-to-charge ratio (*m/z*) of 531was isolated to a purity of 97.5% (Supplementary Fig. 6) and identified it as 5α-cyprinol sulfate (5CS)^23,24^, a known cholanoid (Fig. 2a insets, Supplementary Tables 2, 3 & Figs. 4–13).

**Fig. 2.**
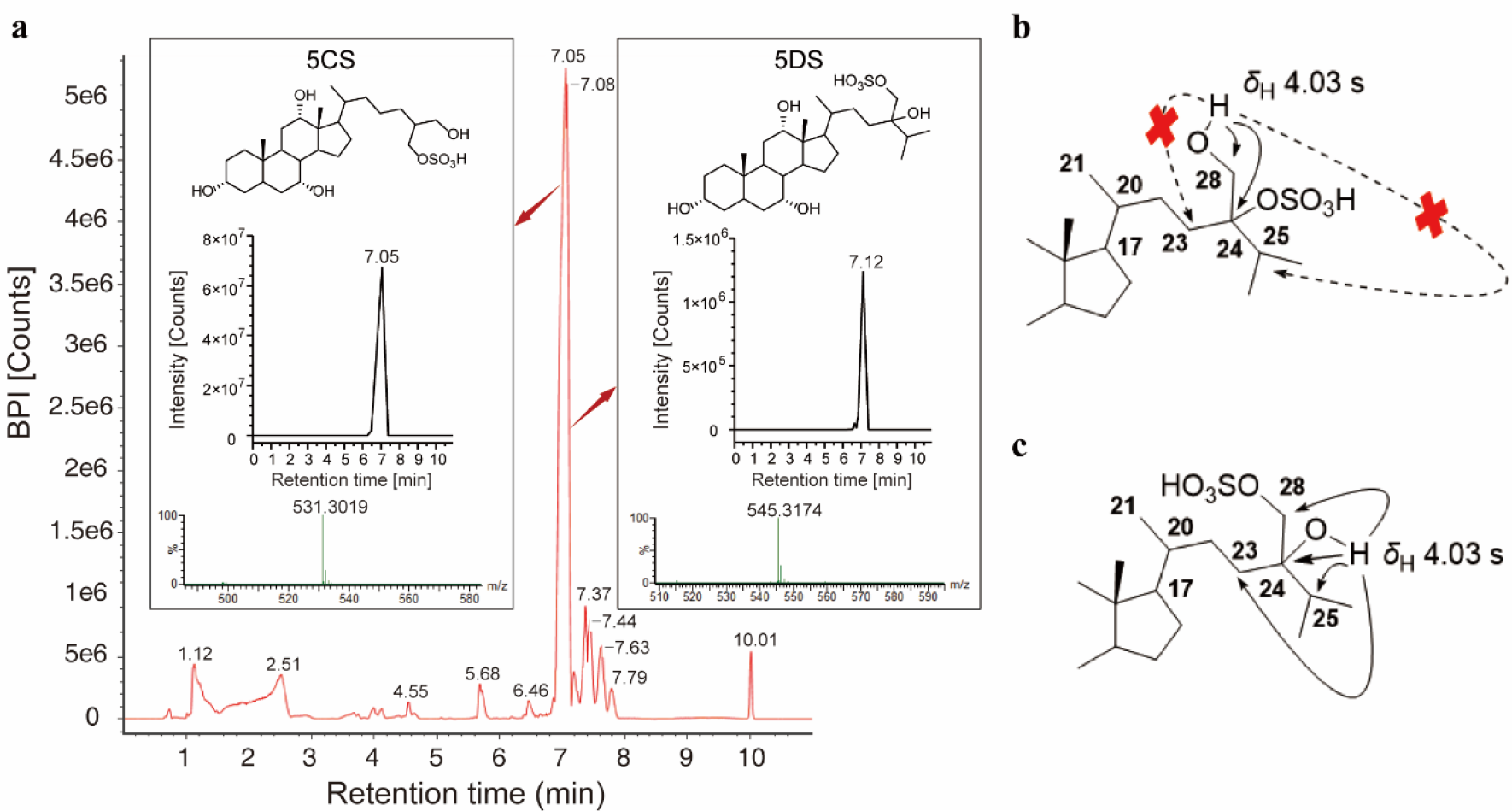
**a)** Chromatography of the n-butanol extract, which shows a predominant peak at retention time 7.05 min that includes 5α-cyprinol (5CS) and 5α-daniol sulfate (5DS) (insets, elucidated chemical structure, ion chromatogram, and mass spectrum, respectively). BPI, base peak ion; *m/z*, mass-to-charge ratio. **b**) and **c**) Key HMBC correlations (arrows) of putative analogs of 5DS. The dashed arrows represent unreasonable correlations.

The compound with *m/z* of 545 was obtained with a purity of 99.1% (Supplementary Fig. 16). The molecular formula was determined to be C_28_H_50_O_8_S based on the HRMS data (*m/z* 545.3149, [M – H]^−^) and the carbon count observed in the ^13^C NMR data (Supplementary Fig. 16 and 18, respectively). This formula represents four degrees of unsaturation. The constitution of the compound was determined to be an oxysterol with a sulfate half ester (−OSO_3_Na) based on the analysis of 1D and 2D-NMR data. The connectivity of the tetracyclic steroidal backbone was determined through analyses of ^1^H–^1^H COSY and HMBC correlations (Supplementary Fig. 14). The ^13^C NMR spectra exhibited distinct resonances corresponding to a 3α,7α,12α-trihydroxy-5α-cholane structure (Supplementary Figs. 18–19). These results suggested that the compound and 5CS shared a tetracyclic structure.

No evidence was observed for additional oxidation in the tetracyclic steroid framework of the molecule with *m/z* 545 in the NMR spectra. Thus, we assumed that the side chain at C-17 consisted of a C_9_H_19_O_5_S subunit. Three methyl groups were identified, each with a single neighboring proton, based on the presence of three three-proton doublets with a coupling constant (*J*) ranging from 6.5 to 6.9 Hz. The COSY and HMBC correlations provided evidence for the presence of a methyl branch at C-20 and C-25, as well as an oxygenated methylene group attached to C-24, which is an oxygenated quaternary carbon (Supplementary Fig. 14). The two-proton doublet (*J* = 2.5 Hz) at 3.62 *ppm* and an oxygenated quaternary carbon at 73.5 *ppm* suggested the presence of a sulfate adjacent to C-24^25^ or C-28^26^ (Figs. 2b–c, respectively). Based on these data, we introduced the side chain as analog B, shown in Fig. 2. Notably, the singlet signal at 4.03 *ppm*, which can be attributed to the hydroxyl either on C-24 or C-28 of the side chain in the COSY spectrum (Supplementary Fig. 20), displayed long-range correlations to C-23 and C-25 in the HMBC spectra (Supplementary Figs. 22–23). These spectra strongly supported the hydroxy substituent at C-25 and sulfation on C-28, respectively. This side chain subunit is present in a sulfated polyhydroxysteroid isolated from starfish, which possesses a different four-ring core backbone^26^. All side chain ^1^H and ^13^C NMR shift data strongly support this constitutional assignment.

The configuration of C-24 could not be assigned because of the high degree of similarity of both the proton and carbon NMR spectra between the C-24*R* and C-24*S* isomers^26^. We are attempting to give an unambiguous answer by using the Mosher reaction when additional quantities become available in the future. The structure was assigned as 24-methyl-5α-cholestane-3α,7α,12α,24,28-pentahydroxy-28-sulfate (Supplementary Fig. 14–15), a novel oxysterol. Herein, we name this compound 5α-daniol sulfate (5DS), acknowledging its origin from *Danio* and the presence of a sulfate ester.

### Mixtures of 5CS and 5DS induced antipredator behaviors

If 5CS and 5DS were alarm substance components, they should, at the concentrations found in 1000×SKE, replicate 1000×SKE in inducing changes in all four behavior parameters. We developed and validated a UPLC-MS/MS method to quantify 5CS and 5DS (Supplementary Figs. 25–26) in SKE (Supplementary Tables 5–7). With this method, we found that the average concentrations of 5CS and 5DS were 1.01×10^−7^ mol/mL and 1.02×10^−9^ mol/mL in 1000×SKE stock solution, respectively (Supplementary Table 8). Based on the diluting procedure in the behavioral bioassay, after adding 1 mL of 1000×SKE stock solution to 1 L of water (Supplementary Fig. 2), the final concentrations of 5CS and 5DS in the 1000×SKE treatment were 1.01×10^−7^ M and 1.02×10^−9^ M, respectively, in the test tank. The 5CS concentrations are approximately 100 (99.19 ± 2.3) times those of 5DS.

We measured zebrafish behavioral responses to 5CS or 5DS, each at concentrations ranging from 10^−6^ to 10^−12^ M. When exposed to 5CS, the zebrafish showed a sharp rise in erratic movement duration at a concentration of 10^−10^ M (Fig. 3a). The time spent in the bottom half showed a significant increase at 10^−12^ M (Fig. 3b). The freezing duration showed no significant changes compared to the vehicle (Fig. 3c). Latency to upper half was elevated remarkably at 10^−10^ M of 5CS (Fig. 3d). Although there was no change in the freezing behavioral phenotype, we tentatively concluded that the threshold for 5CS to induce a partial anti-predator response was 10^−10^ M. After being exposed to 5DS, zebrafish showed increases in all four behavioral parameters at 10^−12^ M (Fig. 3a–d). Thus, the threshold for 5DS to induce an anti-predator response was 10^−12^ M or lower. The locomotion traces showed that 5CS caused more erratic movements (Figs. 3a&i), while 5DS is more likely to trigger freezing (Figs. 3c&j), indicating that distinct components of the alarm pheromone are associated with different behavioral characteristics.

**Fig. 3.**
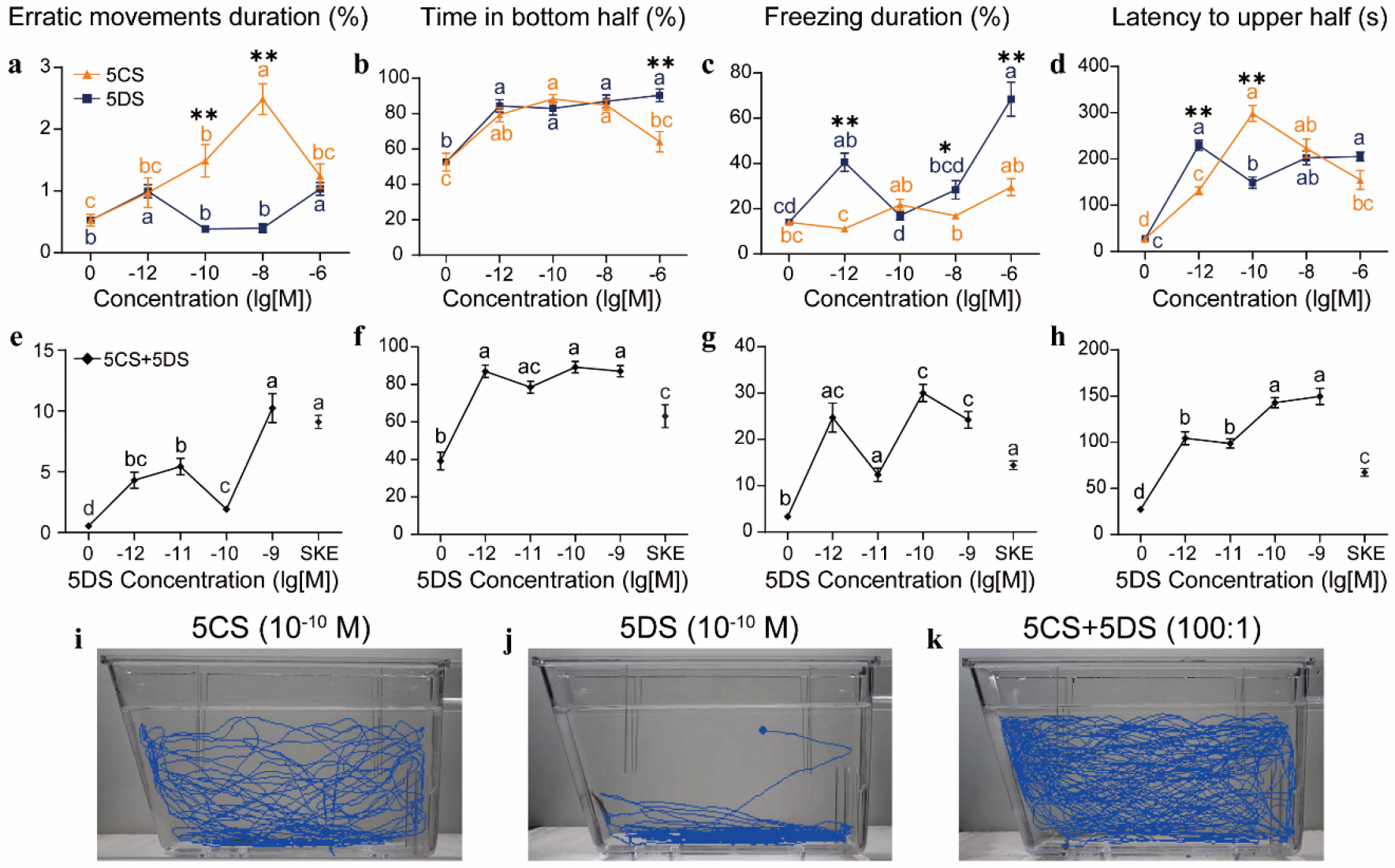
5α-cyprinol (5CS) and 5α-daniol sulfate (5DS) each induced certain anti-predator behaviors: **a**) erratic movement duration (%), **b**) time in the bottom half (%), **c**) freezing duration (%), and **d**) latency to the upper half. The mixture of 5CS and 5DS at a ratio of 100:1 induced all four behaviors (**e**-**h;** 5DS concentrations labeled on the X-axis; 5CS concentrations are 100 times those of 5DS). Skin extract (SKE) was tested at a 1000-fold dilution as a positive control. Representative female motion traces triggered by **i**) 10^−10^ M of 5CS, **i**) 10^−10^ M of 5DS, and **k**) a mixture of 5CS (10^−8^ M) and 5DS (10^−10^ M). * *P* < 0.05, ** *P* < 0.01. Letters above the line indicate significant differences between groups and * or ** indicates a difference in behaviors induced by different compounds at the same concentration. The exact sizes of biologically independent samples and results of ANOVA followed by Bonferroni multiple comparisons ((**a**) two-sided Kruskal–Wallis test (**b** and **f**) and Tamhane test (**c, d, e, g**, and **h**)) are summarized in Supplementary Table 9. In addition, statistical results of behavior comparison caused by different compounds at a specific concentration are summarized in Supplementary Table 10 with independent samples t-test (**a, c**, and **d**) and Mann-Whitney U test (**b**).

### 5CS and 5DS elevated cortisol levels

We further assessed whether 5CS or 5DS induce physiological changes characteristic of zebrafish exposed to alarm or disturbance cues^17^, as in these fish endocrine responses often correlate with behavioral endpoints^27^. Upon exposure to alarm substances^28^ or washings of conspecific individuals that had spotted predators^29^, zebrafish show a rise in cortisol levels. We sacrificed zebrafish immediately after they were recorded for behavioral responses to 5CS and 5DS and prepared samples for cortisol analyses^30^. A significant elevation of cortisol occurred upon exposure to 5CS and 5DS (Fig. 4) at concentrations of 10^−10^ M and 10^−8^ M, respectively.

**Fig. 4.**
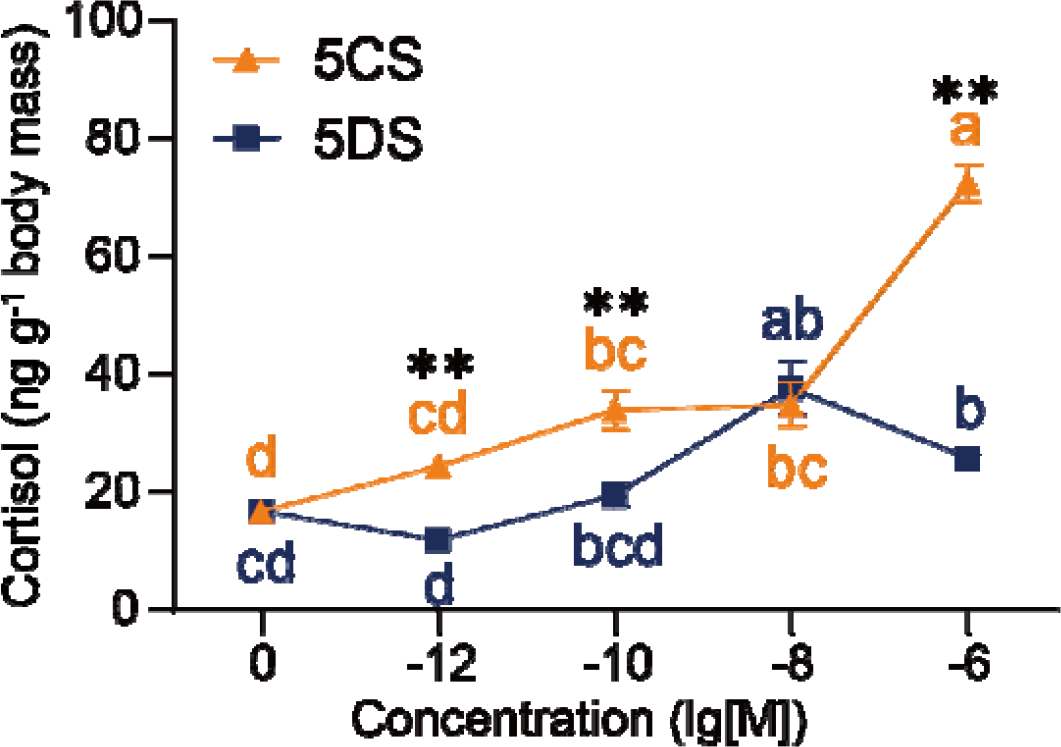
Whole-body cortisol level (ng g^−1^ body mass) in adult zebrafish exposed to 5CS and 5DS at concentrations from 10^−12^ M to 10^−6^ M. Vertical bars, one SEM. For each compound, responses that do not share a letter are different. Asterisk indicates different cortisol levels induced by different compounds at the same concentration (* *p* < 0.05; ** *p* < 0.01). The exact sizes of biologically independent samples and the results of the two-sided Kruskal–Wallis test followed by a multiple comparison of mean ranks are summarized in Supplementary Table 11. Independent samples t-test (concentration (lg[M]) = −12, −10 and −6) and Mann-Whitney U test (concentration (lg[M]) = −8) of the comparison at the same concentration are summarized in Supplementary Table 12.

## Discussion

Our experimental evidence indicates that 5CS and 5DS compose the alarm substance of zebrafish. These two compounds resulted from a stepwise fractionation where the active fraction from each step replicated the skin extract in inducing the conspicuous anti-predatory behaviors. As expected, 5CS and 5DS each induced components of anti-predator behaviors and an elevation of cortisol levels in zebrafish. Of note, individual components of alarm odorants often do not elicit the full behavioral response^16^. The 5CS and 5DS mixture, at the concentrations and ratio found in the skin extract, induced the same set of behaviors as the skin extract. The compounds were effective in stimulating behavioral and physiological responses at sub-nanomolar concentrations. These results demonstrate that 5CS and 5DS are components of zebrafish alarm substance, and are characteristic of the fear substance, as initially described by von Frisch in European minnow^5^ and further defined based on a plethora of chemical ecology studies in teleost fishes^1,2,8,9^.

We designed our bioassay to measure innate behavioral responses and used each test subject only once in the behavioral experiments. The fright reaction in fish is thought to be innate, although prey fish are known to learn to associate odorants with predation or risk of predation^1,8^. We assessed the innate response to reduce variations that could be introduced by associative learning, allowing a consistent measurement of behavioral activities across fractions throughout the stepwise fractionation process. The test was based on the unconditioned model novel tank diving paradigm, which utilizes the animal’s locomotion activity in an unfamiliar setting to assess the fear and anxiety response caused by alarm compounds or synthetic chemicals^15,17^. The behavior phenotypes (e.g., freezing, bottom-dwelling, and erratic movement) and total locomotion are frequently examined in the model and thought to indicate anxiety, fear, and stress^31^. Our findings demonstrated that zebrafish exposed to skin extract displayed increased bottom-dwelling behavior, reduced exploration of the upper region of the tank, and showed more frequent freezing and erratic movement. Compounds 5CS and 5DS, identified using this bioassay-guided approach, replicated the behavioral effects of skin extracts. The behavioral phenotype indicators observed in our experiments were in line with previous findings^14,15,32^.

Our results implicate an additional function of oxysterols, a large family of cholesterol metabolites with broad tissue distribution and physiological functions in animals^33^. Oxysterols have been found in human^34^ and rat^35^ skins and zebrafish embryos^36^. Oxysterol sulfation derivatives 5CS and 5DS are likely to partition into cell membranes like cholesterol^37^ because oxysterols move more rapidly between intracellular membrane organelles due to the greatly increased hydrophilicity of additional hydroxyls and sulfate^38^. 5CS is surface active, increasing the permeability of water-soluble compounds across the mucosal membrane^39,40^. We anticipate that 5CS also has hydrophilic-lipophilic properties because of its similar structure to 5CS. These chemical characteristics enable 5CS and 5DS to exude from injured skin and disperse rapidly in water. Notably, known insect alarm pheromones are thought to dissipate rapidly, which is a characteristic essential to efficient warning of impending danger^4^. As alarm substance components, 5CS and 5DS have the advantage of being released from any injured tissues, including blood, which is also known to contain alarm substances^16^. Moreover, 5CS and 5DS can also be considered derivatives of bile alcohols, a group of metabolites that mediate intraspecific communication and regulate a variety of behaviors in fish, such as foraging, mating, homing, spawning, and migrating^41^. Sulfation is a critical conjugation reaction of bile acid and alcohol, which substantially increases their water solubility and facilitates accessibility to fish’s olfactory organs^42^.

Both 5CS and 5DS appear to possess the biological and chemical features expected of fish alarm subtances^2,43,44^. Because the substances are not species-specific and often elicit equivalent alarm responses in other species, they are postulated to be “public information cues” to which behavior responses have evolved^8^. The survival benefits of these responses have been confirmed in numerous species^1^. Zebrafish are known to shoal with other Cyprinidae fish in their native habitats^45^ and are highly social with other *Danio* species in laboratory culture systems^46^. 5CS is the dominant bile alcohol sulfate of cypriniform fish (including zebrafish)^47^, and is likely released from injured individuals of this taxonomic group, which is known to use alarm substances. Interestingly, 5CS has been shown to act as a kairomone that induces diel vertical migration and the morphological defenses of Daphnia to avoid predators^48,49^. Likewise, chondroitin sulfates derived from tissues of sharks, sturgeon, and *C. elegans* all induced weak fear responses in zebrafish^13^. 5DS has the same bile salt backbone, albeit with an unprecedented adduct of sulfated methylene on C-24. As a novel compound, the biological function and presence of 5DS in other fish species remain to be discovered. If confirmed to be alarm cues in other cypriniform species, 5CS, and possibly 5DS, could become molecular templates to study how related species respond to shared alarm cues. Alarm substances, sometimes referred to as alarm pheromones, are likely less specialized than other types of pheromones^4^, which may be useful for fish that shoal together to warn and detect shared predation threats. It would be interesting to determine if 5CS or 5DS (or both) induces anti-predation behaviors in other species, particularly those that shoal with zebrafish. Such studies may provide useful information to infer how alarm substances evolved as a result of the evolutionary arms race between prey and predators.

The alarm substance components identified in this study provide a molecular model to examine the broad function of danger signaling in fish. The zebrafish is extensively used as a model organism in studies of behavior^50^, physiology^51^, and development^52^. In this species, the alarm substance elicits innate fear, manifested by changes in conspicuous behaviors^15,17^, anxiety levels^15,17^, and early development^53^. Zebrafish embryos are known to alter the development progress when exposed to adult alarm substances^53^, while crushed embryos release alarm substances that induce anti-predator behaviors in adults^54^. 5CS and 5DS would enable us to interrogate how the olfactory receptor neurons and the central nervous system mediate responses to danger and incite fear throughout life history, which will further support the utility of zebrafish as a model animal in studies related to human anxiety^55,56^.

## Online content

Experimental material and methods, Result details, and Spectra for elucidation of compounds 5CS and 5DS are available in Supplementary Materials.

## Supporting information

Movie of erratic movement

Movie of freezing

## Acknowledgments

We thank Ying Kong in Yantai Branch, Shanghai Institute of Materia Medica, Chinese Academy of Sciences, for assistance with quantitative analysis method development and validation for 5CS and 5DS.

## Funding

This study received funding from the National Natural Science Foundation of China (32270533). K. L. expresses gratitude for the Taishan Scholar Program provided by the Shandong Province of China (tsqn20190403), as well as the support received from the Yantai Municipal City’s Shuangbai Plan (2018020). W. L. is supported by the Great Lakes Fishery Commission.

## Author contributions

The study was designed by K. L and W. L. Y. L. and Z. Y. were responsible for maintaining the aquarium. Y. L., Z. Y., X. Y., and X. C. conducted extractions and behavioral analysis. A. L collected the skin. Y. L., Z. Y., K. L., X. L., and X. Y. conducted structure elucidation and quantitative analysis. Y. L did the statistical analysis. K. L. obtained funding. The manuscript was written by K. L. and W. L. with contributions from all authors. K. L. provided supervision for the project.

## Competing interests

The authors declare no competing interests.

## Materials and Methods

### Zebrafish housing

All experiments were performed following the guidelines of the Animal Care Ethics Committee of the Yantai Institute of Coastal Zone Research, Chinese Academy of Sciences (2021R001). Adult zebrafish (3–5 months old; average weight, 0.2 g; average body length, 3.1 cm) were acquired from a commercial distributor (Qingfeng Fisheries, Shanghai, China). The fish were housed in a recirculating system (Haisheng, Shanghai, China), with water that had been treated through a water filtration system (Haier, Qingdao, China) and supplemented with sea salt to maintain a concentration of approximately 750 ppm. Each tank (10 liters) contained 10-20 fish. The water temperatures were kept within the range of 25–27°C. Ceiling-mounted fluorescent light tubes were used for illumination on a 12-hour cycle, with the lights turning on at 08:00 and turning off at 20:00. Adult zebrafish were fed daily with brine shrimp (BESSN, Beijing, China).

### Behavioral bioassay

The behavioral bioassay, a novel tank diving test, was modified from Cachat *et al*.^15^. The experiment was conducted using a trapezoidal prism aquarium tank with dimensions of 27×7 cm (upper), 22×7 cm (lower), and a depth of 15 cm. This tank, referred to as a “novel tank,” was supplemented by a pretreatment beaker with a capacity of 1 L (Fig. S1). Zebrafish were initially exposed to alarm substances for 5 minutes^17^ in the pretreatment beaker before being moved to a novel tank for behavioral observation and recording. The control group underwent the same procedure and was exposed to the appropriate vehicle stimulus for each experiment. The fish behaviors were documented using a Canon EOS Rebel T3 camera immediately when the fish was transferred into the novel tank. The recording lasted for 7 minutes. The fish motion was tracked using an EthoVision XT10 (Noldus Information Technology, Wageningen, the Netherlands) based on recorded videos. The tracking analysis commenced once the subject fish was detected for a duration exceeding 1 s. The detection settings were chosen to precisely capture the alarm behavior of zebrafish^15^. The movement tracks were smoothed and analyzed for abnormalities using EthoVision XT10. Next, a standard two-dimensional swim track was generated.

### Analysis of behavioral data

The behavioral data of zebrafish were analyzed using computer-aided analysis. Two trained observers conducted the analysis using a double-blind procedure. We evaluated 12 behavioral parameters: time spent in the bottom (%), latency to enter the top (s), freezing time (%), duration of erratic movements (%), number of entries to the bottom, distance traveled in the bottom, average velocity, total distance traveled, time spent in the bottom and top, entries in the bottom and top, average duration of bottom entries (s), and distance traveled in the bottom and top. Commercial software offered an efficient method for quantifying behavioral endpoints, including time spent at the bottom, latency to enter the top, number of entries to the bottom or top, distance traveled, and average velocity. In addition, manual analysis was conducted to examine freezing and erratic movements^15^. In this study, freezing refers to a state of immobility in fish, characterized by limited movement in the gills and the eyes. This behavior is predominantly observed when the fish is positioned near the bottom, in a corner, or just beneath the water’s surface^57^. The erratic movement referred to is characterized by a high swim speed of more than 3 cm/s and a seemingly aimless zig-zag pattern with frequent changes in swimming direction^57^. This behavior is commonly observed at the bottom of the tank, but can also occur in midwater. The remaining parameters can be quantified through computerized video tracking and automatic recording, and subsequently determined using the formula^58^. The data were plotted using GraphPad Prism software (GraphPad, NJ, USA) and statistically analyzed using SPSS (version 11.0, SPSS Inc, IL, USA).

### Extraction, fractionation, and bioassay of compounds from the skin

The zebrafish (approximately 800 fish) were caught using a net and anesthetized with tricaine spray (Sigma-Aldrich, St. Louis, MO, USA) on an ice plate. The skin of each fish was peeled off using dissecting scissors and forceps on the ice plate. The harvested skin samples (46.2 g) were homogenized (60 Hz, 90 s) and extracted by ethanol three times (95%, 3 × 300 mL, vortex 10 min each, Merck, Germany). The three batches of extracts were pooled together and centrifuged for 20 min at 10000*g* and 4°C (Bruker, Rheinstetten, Germany). The supernatants were concentrated using a roto-evaporator (model R-210, Buchi Rotavapor, Flawil, Switzerland). The residue (skin extract; SKE, *ca*. g) was separated into three parts, with 5% (0.38 g) of the residue used for behavioral assays, 80% (6.0 g) used for compound isolation, and the remaining 15% (1.1 g) archived (Fig. S2).

We tracked the alarm substance activities in each fractionation step and used the results to guide our search for the active compounds (Fig. S2). The residue (5%) was dissolved in 0.5 mL of water to acquire a stock solution equivalent to 80 fish/mL (designated as SKE stock solution), diluted serially from 10 to 10^6^-fold, resulting in “stimulus solutions”. Each stimulus solution was assayed by adding 1 mL of stimulus solution to the 1 L pretreatment beaker (Fig. S1). The 10 to 10^6^ fold stimulus solutions in the pretreatment beaker corresponded to 8 fish extract/L to 8×10^−5^ fish extract/L (Fig S2). This dilution scheme is similar to the alarm substance extract protocol^14,19,20^ and the odorant concentration applied in alarm response experiments (1×10^−3^ fish extract/L)^21^.

Subsequently, a portion of the residue (80%; 6.0 g) was suspended in 100 mL of water and subjected to sequential partitioning with petroleum ether (PET), ethyl acetate (EtOAc), and n-butanol (nBuOH) (3 × 100 mL each, Merck, China). The extract of each phase was concentrated in *vacuo*, resulting in extracts of PET (1.2 g), EtOAc (0.3 g), and nBuOH (4.1 g). These fractions of skin extract, namely, PET, EtOAc, and nBuOH, were each reconstituted into 8 mL of water (Fig. S2). From each fraction, we took 0.1 mL from the 8 mL stock solution and mixed it with 99.9 mL water (1000 fold dilution), resulting in a test solution equivalent to 1000×SKE (0.08 fish/mL) and subjected these solutions to the behavioral bioassay (Fig. S2). The test solutions were assayed as described for SKE stimulus solutions. The final treatment concentration in the baker was 0.08 fish/L. The behavioral outcomes were compared to those elicited by the 0.08 fish extract (1000×SKE treatment) to determine which fraction or fractions contained alarm substance components.

The n-butanol test solution replicated the potency of 1000×SKE (0.08 fish/mL) in the bioassay. Hence, the residue of nBuOH extract (4.05 g, 98.7% of n-butanol extract) was chromatographed by high-performance liquid chromatography (HPLC) coupled with a quaternary gradient pump, a photodiode array detector (Waters, Milford, MA, USA). We used a reverse phase column (Waters Xbridge, BEH C18 10 × 250 mm, 5 μm) at room temperature and eluted fractions with a mobile phase consisting of water (A) and methanol (B) at 4.0 mL/min. Separation was achieved using the following gradient program for 50 min: 95% A for 5 min, decreased to 5% A from 5 to 40 min, and then maintained at 5% A from 40.0 to 50.0 min. The manual injection volume was 1 mL. The five-minute eluents contained a large amount of water and methanol and were lyophilized. Fraction 7 was reconstituted to 7.9 mL (Fig. S2), from which 0.1 mL was taken and diluted with 99.9 mL of water, resulting in a 10^3^-fold diluted solution that was equivalent to 0.08 fish/mL (1000×Fr.7), and 1 mL of this solution was added into 1 L of water in the pretreatment beaker and proceeded to be assessed for its behavioral activities.

Fraction 7 (the remaining 7.8 mL) was concentrated to obtain a residue (24 mg) that was further purified using Sephadex LH-20 (Sigma-Aldrich, Shanghai, China), eluted first by CH_2_Cl_2_-MeOH (1:1) and then by MeOH (100%), which resulted in compounds **1** (7.8 mg, 5CS) and **2** (1.8 mg, 5DS).

Compound 5CS (1 mg) was reconstituted in 1.8 mL of water to make a 10^−3^ M stock solution and, subsequently, diluted serially to 10^−9^ M. Similarly, 1 mg of 5DS was reconstituted in water to form 1.8 mL of 10^−3^ M stock solution and diluted to prepare a treatment solution ranging from 10^−6^ to 10^−9^ M (Fig. S2). We also combined 100 μL of 5CS (10^−3^ M) and 100 μL of 5DS (10^−5^ M) and diluted this combined solution to 1 mL to make a stock solution of the mixture (5CS, 10^−4^ M, and 5DS, 10^−6^ M). The solution was serially diluted. The novel tank diving tests were carried out on 5CS, 5DS, and the mixture, respectively. In each trial, 1 mL of test solution was dissolved in 1 L of water to prepare a treatment with concentrations as shown in Fig. S2.

### NMR analysis

The NMR spectra of the compounds were obtained at 23°C using a Bruker Avance III 500 NMR spectrometer with a 5 mm tube. The spectrometer operated at 500 MHz for ^1^H and 125 MHz for ^13^C. The ^13^C DEPT spectra (135°, 90°, and 45°) were employed to distinguish between the resonance signals of methine and methylene groups. The research employed ^1^H and ^13^C resonances, along with a set of two-dimensional (2D) homonuclear (^1^H-1H) and heteronuclear (^1^H-^13^C) shift-correlated approaches. The employed methodologies encompassed ^1^H-detected heteronuclear multiple quantum correlation (HMQC), ^1^H-detected heteronuclear multiple bond correlation (HMBC), ^1^H-^1^H correlation spectroscopy (COSY), and ^1^H-^1^H nuclear Overhauser effect spectroscopy (NOESY) experiments. The 2D-NMR spectra were obtained using recommended pulse sequences and parameters provided by the manufacturer.

### Quantitative analysis of 5CS and 5DS

We developed and validated a LC-MS/MS method to quantify 5α-cyprinol sulfate (5CS) and 5α-daniol sulfate (5DS), in zebrafish skin. Quantitative analysis was conducted using a UPLC system (UltiMate 3000, Thermo Fisher Scientific, Waltham, MA, USA) coupled with a TSQ Quantiva mass spectrometer (Thermo Fisher Scientific). A Thermo Hypersil Gold C18 column with dimensions of 2.1 × 50 mm and particle size of 1.9 μm was employed for the separation process. The mobile phase comprised solvent A, which was water containing 0.1% formic acid (FA), and solvent B, which was methanol. Gradient elution was performed using the following conditions: 0–0.2 min with 10% B, 0.21–2.0 min with a linear increase from 10% B to 90% B, 2.0–4.0 min with 90% B, 4.01–4.5 min with a linear decrease from 90% B to 10% B, and finally, the column was re-equilibrated with 10% B for 1.0 min. The flow rate was 0.3 mL/min. The column temperature was set to 40 °C, and the sample injection volume was 1.0 µL. The autosampler syringe was washed with methanol during the intervals between injections.

The TSQ Quantiva was operated in negative mode. The selected reaction monitoring transitions for determination were monitored under optimal conditions. The monitoring ion pairs used for 5CS and 5DS were 531.3→97.0 and 545.3→97.0, respectively. The collision energies for 5CS and 5DS were optimized to 55 V, while the RF lens was set at 190 V for both analytes. The following mass parameters were configured: negative ion at 2500 V, sheath gas at 35 arb, auxiliary gas at 15 arb, sweep gas at 2 arb, ion transfer tube temperature at 350 °C, vaporizer temperature at 300 °C, and CID gas at 1.5 mTorr. The dwell times for both analytes were 100 ms. The Xcalibur software, developed by Thermo Fisher Scientific, controlled the MS system. The 1000×SKE solution was diluted and analyzed using UPLC-MS/MS to quantify the absolute concentration of 5CS and 5DS (Fig. S2).

### Quantitative analysis of cortisol

The method developed and validated by our laboratory was used to extract and determine the whole-body cortisol of zebrafish^59^. Briefly, the fish were euthanized using tricaine at a concentration of 500 mg/L in the ice bath, removed of excessive water using paper towels, frozen in liquid nitrogen, and stored at –80°C. to process the samples, the fish were thawed, weighed, diced, and then homogenized at 60 Hz for 90 seconds. The sample was subsequently centrifuged at 10,000 *g* and 4°C for 10 minutes. The stable isotope internal standard cortisol-*d*_4_ (500 ng, Sigma-Aldrich Corp.) was added to the fish supernatant. Solid-phase extraction (SPE) with the Waters Oasis HLB Vac Cartridge that had been pre-conditioned with 10 mL of methanol and 10 mL of water was used to extract the cortisol from the supernatant. Samples were processed using a 1 mL/min Supelco vacuum manifold (Sigma-Aldrich Corp.), which can extract 12 samples concurrently. To reduce interference, loaded cartridges were rinsed with a solution consisting of 30% methanol/water and 2% acetic acid (5 mL). Subsequently, the cartridges were washed with a solution of 75% methanol and 2% acetic acid in water (5 mL). Eluates were collected and concentrated using a Cold Trap with Vacuum Centrifuge Concentrator (JM Technology Co., Beijing, China). The concentrated eluates were then reconstituted in 100 μL of methanol/H_2_O (55:45, v:v) for UPLC-MS analysis. The UPLC-MS method validated following FDA guidelines was used to accurately measure cortisol concentration in whole-body homogenates of zebrafish^59^.

### Statistics

All behavior and endocrine data were tested for normality. For sets of data that conformed to a normal distribution, an independent samples t-test (for data from two treatments) or one-way ANOVA (for data from three or more treatments) was used, with Bonferroni correction where needed.

To define the parameters that describe the full spectrum of behaviors induced by all components of AS, or SKE, data from all 12 measured behavior parameters were pooled and analyzed. The data did not conform to normality, and various transformations did not correct the issue. We, therefore, decided not to perform a MANOVA on the pooled data. Data from each of the 12 parameters were subjected to independent sample t-tests if the data conformed to the normal distribution and the Mann-Whitney U test if the data did not conform to the normal distribution. All tests were subjected to Bonferroni correction (with total tests = 12).

Bioassay data from the products of each fractionation step were treated as the results of an independent experiment. To account for possible block effects due to the lengthy process of fractionation and the different solvents used, blank control tests using appropriate vehicle stimuli were performed for each set of experiments. The four parameters determined to represent anti-predation behavior were analyzed with either an independent sample t-test or a Mann-Whitney U test, with Bonferroni correction. For data that do not conform to a normal distribution, a non-parametric test is used. For each assay, tests were repeated ten times (n = 10). A *p*-value□<□0.05 was considered to indicate significance.

Bioassay data for purified compounds were also analyzed with one ANOVA or Kruskal-Wallis test, and with *post hoc* tests. Cortisol data were analyzed similarly.

## Notes

### Competing Interest Statement

The authors have declared no competing interest.

