## Supplementary material for "Novel Epidermal Oxysterols Function as Alarm Substances in Zebrafish"

**This PDF file includes:**

Supplementary Text
Figs. S1 to S26
Tables S1 to S13

### Supplementary Text

#### ***Chemical structure elucidation of 5CS***

Compound **5CS** was obtained as a pale amorphous powder. The high resolution negative ion mass spectral data gave a pseudomolecular ion of *m/z* 531.3019 $[M - H]^-$, corresponding to the molecular formula of $C_{27}H_{48}O_8S$, (cal. 531.2992 $[M - H]^-$, $\Delta = 5.1$ ppm) (Fig. S6). The collision-induced fragmentation of this ion resulted in the detection of a product ion at *m/z* 452, which coincided with that of grass carp bile alcohol[1]; the latter was attributed to the monoisotopic mass of hydrogen sulfate anion ($HSO_4^-$, *m/z* 98). The latter ion indicated the presence of a sulfate group and implied four degrees of unsaturation.

The constitution of **5CS** was determined by 1D- and 2D-NMR analyses (Figs. S7-S13 and Table S2-S3). The $^1$H, $^{13}$C, and HSQC NMR spectra (Figs. S7-S9 and S11) of **5CS** in methanol-$d_4$ showed signals for two methyl singlets representing the methyl groups C-18 and C-19 and a doublet for the methyl groups C-21. These methyl group signals and the predicted molecular formula indicated a classic 27-carbon steroid skeleton. Among the HMBC correlations (Fig. S4) for **5CS** was a long-range communication between the methyl protons for Me-19 to a CH ($\delta_C$ 32.9). The COSY spectrum (Fig. S10) showed correlations between $CH\text{-}CH_2\text{-}CH\text{-}CH_2\text{-}CH$. The proton on C-5 also showed HMBC correlations to the CH at $\delta_C$ 67.4 (Fig. S4) and CH at $\delta_C$ 68.9, indicating the oxygenation on C-3 and C-7. The long-range communication between the methyl protons for Me-18 to carbinol methine proton at $\delta_C$ 74.1 allowed us to locate another site of oxygenation on C-12. These relationships were highly reminiscent of the hydroxylation pattern in petromyzonol [2]. The chemical shifts of all five carbons C-3–C-7 matched well with those of petromyzonol (Table S2).

Accounting for the molecular formula deduced by HR-MS, we assumed the composition of the C17 side chain to be a $C_8H_{17}O_5S$ subunit. A $CH_2$ resonance at $\delta_H$ 4.03 was attached to a carbon having a chemical shift of $\delta_C$ 69.1 (HSQC). This is a characteristic set of chemical shifts for sulfated primary alcohol [3], in addition to another primary alcohol $CH_2$ resonance at $\delta_H$ 3.60 and $\delta_C$ 63.3, which accounted for all heteroatoms present in **5CS**. In the HMBC spectrum (Fig. S12), the $CH_3$ resonance at $\delta_H$ 1.02 showed a long-range correlation with $\delta_C$ 48.5 (C-17), which also exhibited a strong correlation with Me-18. In addition, the COSY correlation between H-21 and H-20 and the correlation between H-20 and H-17 were supportive of the connectivity between the side chain and tetracyclic steroidal backbone. Therefore, the planar structure was assigned as 3,7,12-trihydroxy-cholestan-27 sulfate, consistent with that of cyprinol sulfate [4,5].

The relative configuration of **5CS** was supported by NOESY (Fig. S5) and coupling constant analysis. The NOESY correlations observed from H-19 to H-8$\beta$, from H-8$\beta$ to H-18, from H-18 to H-21, as well as on the counterpart, the correlations from H-5α to H-9α, from H-9$\alpha$ to H-14$\alpha$, and from H-14$\alpha$ to H-17$\alpha$, indicated the relative configuration for each ring junction to be *trans*. Furthermore, the NOESY spectrum (Fig. S13) indicated the following  orrelations: from H-3 to H-1$\beta$ and from H-1$\beta$ to H-19; H-7 with H-8$\beta$; H-12 with H-11$\beta$, which also presented correlation with H-8$\beta$. These features indicated the hydroxyls at C-3, C-7, and C-12 have an axial orientation [4]. The chemical shifts of **5CS** showed similarity to those of 5$\alpha$-cyprinol sulfate and a significant difference from those of the 5$\beta$ analog (Fig. S5 and Table S2 & S3).

In addition, the optical rotation of **5CS** ($[\alpha]^{25}_D$ +38.0, *c* 0.20, MeOH) was consistent with that of isolated 5$\alpha$-cyprinol sulfate [4]. Therefore, we confirmed that the chemical structure is 5$\alpha$-cyprinol sulfate.

***Chemical structure elucidation of 5DS***

Compound **5DS** was obtained as a white amorphous powder. The molecular formula of **5DS** was established as $C_{28}H_{50}O_8S$ from its HRESIMS (*m/z* 545.3149, $[M - H]^-$) (Fig. S16) in conjunction with the carbon count indicated by the $^{13}$C NMR data (Fig. S18). This indicated four degrees of unsaturation. The constitution of **5DS** was deduced to be an oxysterol with a sulfate half ester ($-OSO_3Na$) through analysis of 1D- (Table S4) and 2D-NMR (Fig. S17-S23) data. The connectivity of most of the tetracyclic steroidal backbone was established by interpretation of the $^{1}$H–$^{1}$H COSY and HMBC correlations (Figs. S20&S22-23). The $^{13}$C NMR spectrum showed characteristic resonances for a 3*α*,7*α*,12*α*-trihydroxy-5*α*-cholane skeleton (Fig. S18). These results suggested that **5DS** shared a tetracyclic structure common to that deduced earlier to be present in **5CS**.

There was no evidence in the NMR spectra for additional oxidation in the tetracyclic steroid framework. Therefore, we assumed that the composition of the C17 side chain was a $C_9H_{19}O_5S$ subunit. Three of the three-proton doublets ($J$ = 6.5−6.9 Hz) indicated three methyl groups, each having a single vicinal proton. The COSY and HMBC correlations supported the methyl branching at C20 and C25, and an oxygenated methylene substituent attached to C24, an oxygenated quaternary carbon. The two-proton doublet ($J$ = 2.5) at 3.62 ppm and an oxygenated quaternary carbon at 73.5 ppm suggested the presence of a sulfated adjacent to C24 [6] or C28 [7] (B&C in Fig. 2, respectively). To accommodate these facts, we proposed the side chain as analog B shown in Fig. 2. Notably, the singlet signal at 4.03 ppm, which can be attributed to the hydroxyl either on C-24 or C-28 of the side chain in the COSY spectrum, displayed long-range correlations to C-23 and C-25 in HMBC spectrum which strongly supporting the hydroxy substituent at C-25 and sulfated on C-28, correspondingly. This substructural unit is present in the sulfated polyhydroxy steroid isolated from starfish, which possesses a different steroid backbone[7]. The $^{1}$H and $^{13}$C NMR shift data for all side chain protons strongly support this constitutional assignment for **5DS**.

While it is tempting to propose that **5DS** and reference compound[7] also share the same configuration at C-24, we have been unable to locate any reported examples that are epimeric at C-24 to assess whether chemical shift differences were distinctive. The 24*R* configuration in reference compound [7] was assigned by the enantioselective synthesis of (24*R*-) and (24*S*-)-24-hydroxymethyl-24-hydroxycholesterols. The comparison presented a high degree of similarity of the proton and carbon NMR spectra for each of the C-24-epimers, and an unambiguous answer to this question must await resolution by synthesis, which we are pursuing to solve by Mosher reaction when more amounts of **5DS** become available in future [7].

***Quantitative analysis of 5CS and 5DS in the AS stock solution and skin***

LC-MS/MS has been widely used in the quantification of oxysterols due to its selectivity and specificity. After optimization of chromatographic conditions, the analytes 5CS and 5DS were well separated on a Thermo Hypersil Gold C18 column as shown in Fig. S26. Using the optimized SIM parameters displayed in Table S5, the target analytes in the present study could be quantified reproducibly and accurately.

Linearity was determined by analysis of a 9-point calibration curve. Charcoal-stripped serum after the removal of bile acids was used as the blank serum matrix for the preparation of calibrators. The linear correlation coefficients ($r^2$) of the calibration curves were 0.9992 for 5CS and 0.9996 for 5DS, respectively, suggesting excellent linearity for the quantification. A high level of linearity in responses over a sufficient dynamic concentration range was observed and summarized in Table S6. The intra- and inter-day accuracy and precision were evaluated using 3 QC concentrations distributed throughout the calibration range for each analyte (Table S7). Intra-day accuracy (n = 5) ranged from 100.60 to 105.03%. For precision of analytes, intra-day variation (n = 5) ranged from 1.18% to 4.15%. Inter-day accuracy (n = 3) ranged from 97.59 to 108.03%. For precision of analytes, inter-day variation (n = 3) ranged from 2.51% to 4.32%. The precision and accuracy for all analytes were more than 85%, which was within the interval set by the US Food and Drug Administration (FDA) concerning the validation of bioanalytical methods. Therefore, the developed method yields excellent reliability and reproducibility.

The method has been used for quantitative analysis of 1000×SKE stock solution. The concentrations of 5CS and 5DS in the sample were 535.64 ± 8.33 ng/mL and 5.54 ± 0.14 ng/mL (n = 5), respectively. According to our dilution procedure of concentration determination, we deduced that the concentrations of 5CS and 5DS were $1.03\times10^{-7}$ mol/mL and $1.12\times10^{-9}$ mol/mL. After adding 1 mL to 1 L water, the final concentrations of 5CS and 5DS in 1000×AS treatment were $1.03\times10^{-7}$ M and $1.12\times10^{-9}$ M, respectively. The mixture of $10^{-7}$ M of 5CS and $10^{-9}$ M of 5DS (approximately the same concentrations as those in the 1000×SKE treatment) was prepared with isolated compounds and serially diluted to complete the bioassays.

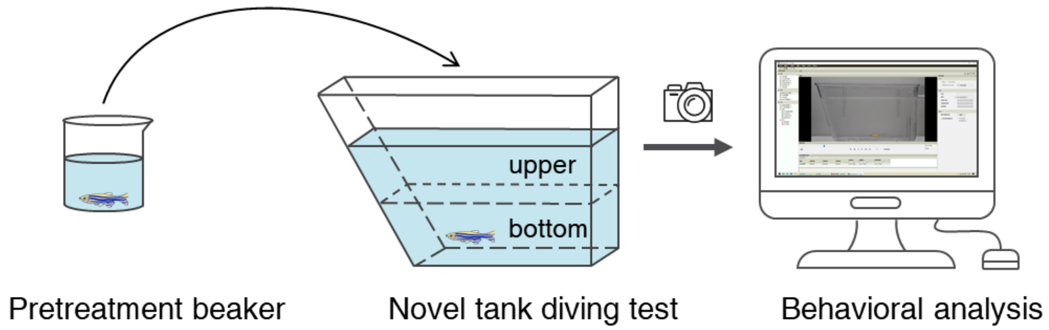


**Fig. S1** Schematic diagram of behavior test and data analysis of zebrafish. A 1.5 L beaker filled with 1L stimulus as pretreatment. The fish was transferred to the novel tank trapezoidal prism aquarium tank (27×7 cm upper, 22×7 lower, 15 cm in depth).

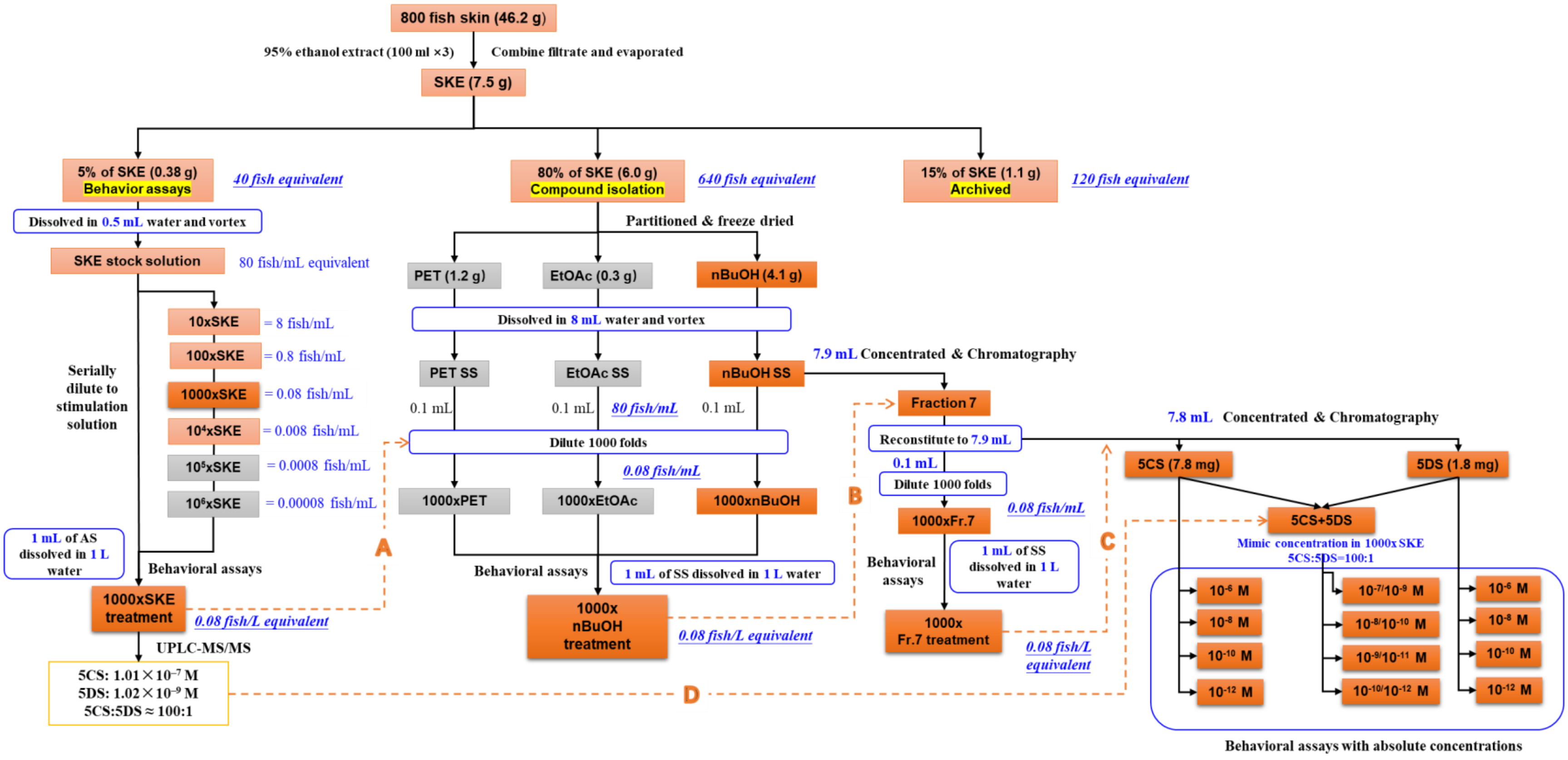


**Fig. S2** Bioassay-guided strategy to isolate active components of alarm substances from zebrafish skin. The orange boxes represent the treatment with behavioral activities; the grey boxes represent inactive fractions; the blue boxes present dilution steps. The black arrows represent the dilution procedure of SKE, successively partition, and the isolation of compounds **5CS** and **5DS**; the orange dashed arrows represent the behavior evidence used to guide the chemical procedure. The test solutions for behavior treatments, 1000×SKE, 1000×nBuOH, and 1000×Fraction7 were all adjusted to be equivalent to the extract of 0.08 fish/L. (**A**) The 1000×SKE treatment showed activity in all four behavioral phenotypes, suggesting diluting 1000 fold for each phase of fractionation was appropriate. (**B**) The 1000×nBuOH treatment replicated all the behavioral activities of 1000×SKE treatment, indicating the active compounds are in nBuOH phase. (**C**) Further fractionation and bioassay indicated fraction 7 replicated the activity of nBuOH phase. Finally, compounds **5CS** and **5DS** were identified from Fraction 7 of the nBuOH phase. (D) The concentration of **5CS** in 1000×SKE treatment water was $1.01\times10^{-7}$

M. Similarly, the concentration of **5DS** in 1000×SKE treatment water was $1.02\times10^{-9}$ M. The mixture of $10^{-7}$ M of **5CS** and $10^{-9}$ M of **5DS** was prepared with isolated compounds and serially diluted to complete the bioassays.

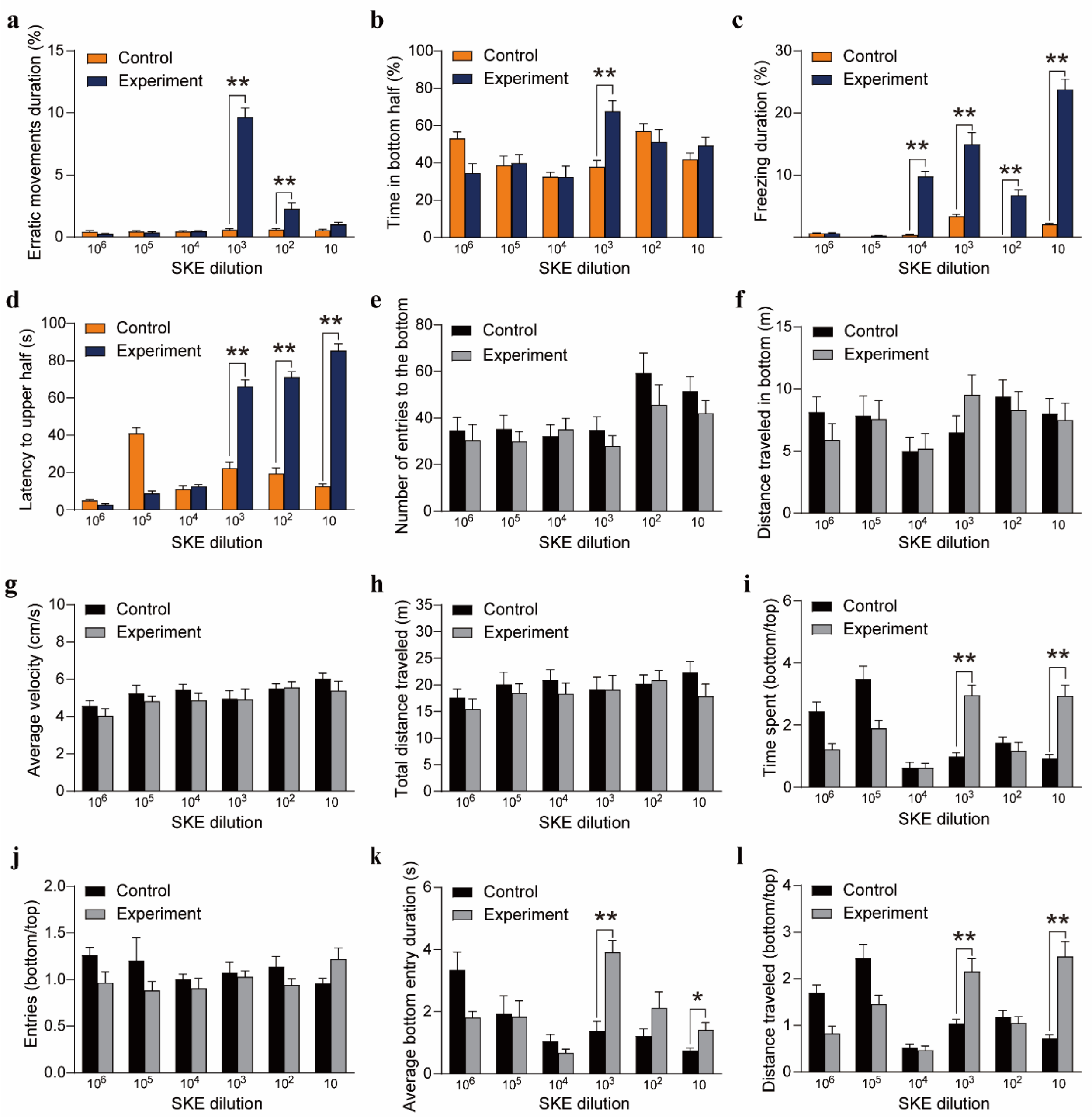


**Fig. S3** Zebrafish anti-predation behavior phenotypes triggered by SKE dilutions, **a**) erratic movements duration (%) **b**) time in bottom half (%), **c**) freezing duration (%), **d**) latency to upper half (s), **e**) number of entries to the bottom, **f**) distance traveled in bottom (m), **g**) average velocity (cm/s), **h**) total distance traveled (m), **i**) time spent (bottom/top), **j**) entries (bottom/top), **k**) average bottom entry duration (s), **l**) distance traveled (bottom/top). Asterisks indicate significant differences between treatment and control (*$P < 0.05$, **$P < 0.01$). Data are means ± SEM. Exact number $N$ of biologically independent samples and results of independent samples t-test or Mann-Whitney U test are summarized in Supplementary Table 13.

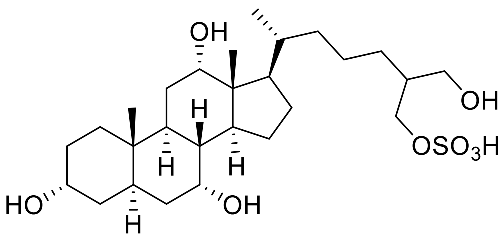


***5CS***

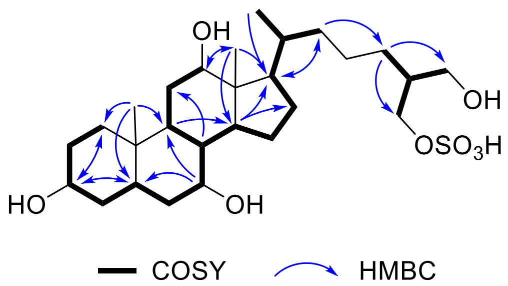


**Fig. S4** Chemical structure and key COSY and HMBC correlations of 5$\alpha$-cyprinol sulfate (**5CS**)

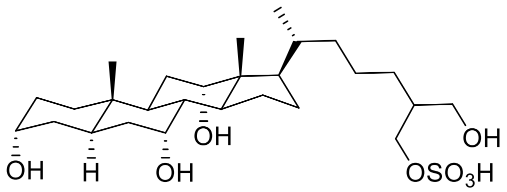

**5*α*-cyprinol sulfate**

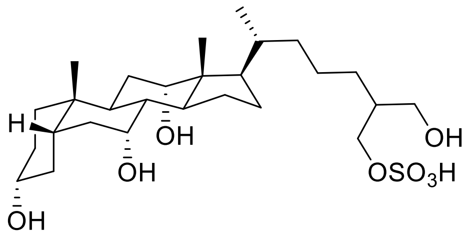

**5*β*-cyprinol sulfate**

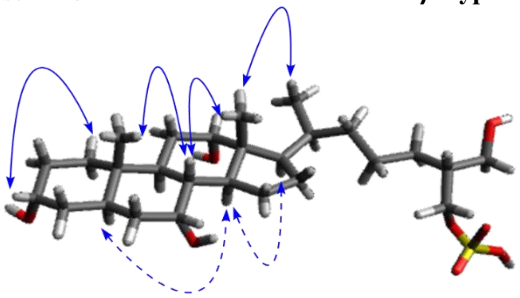

**5CS**

**Fig. S5** Chemical structure of 5*α*-cyprinol sulfate (**5CS**) and its epimer 5*β*-cyprinol sulfate, and NOESY correlations of **5CS**

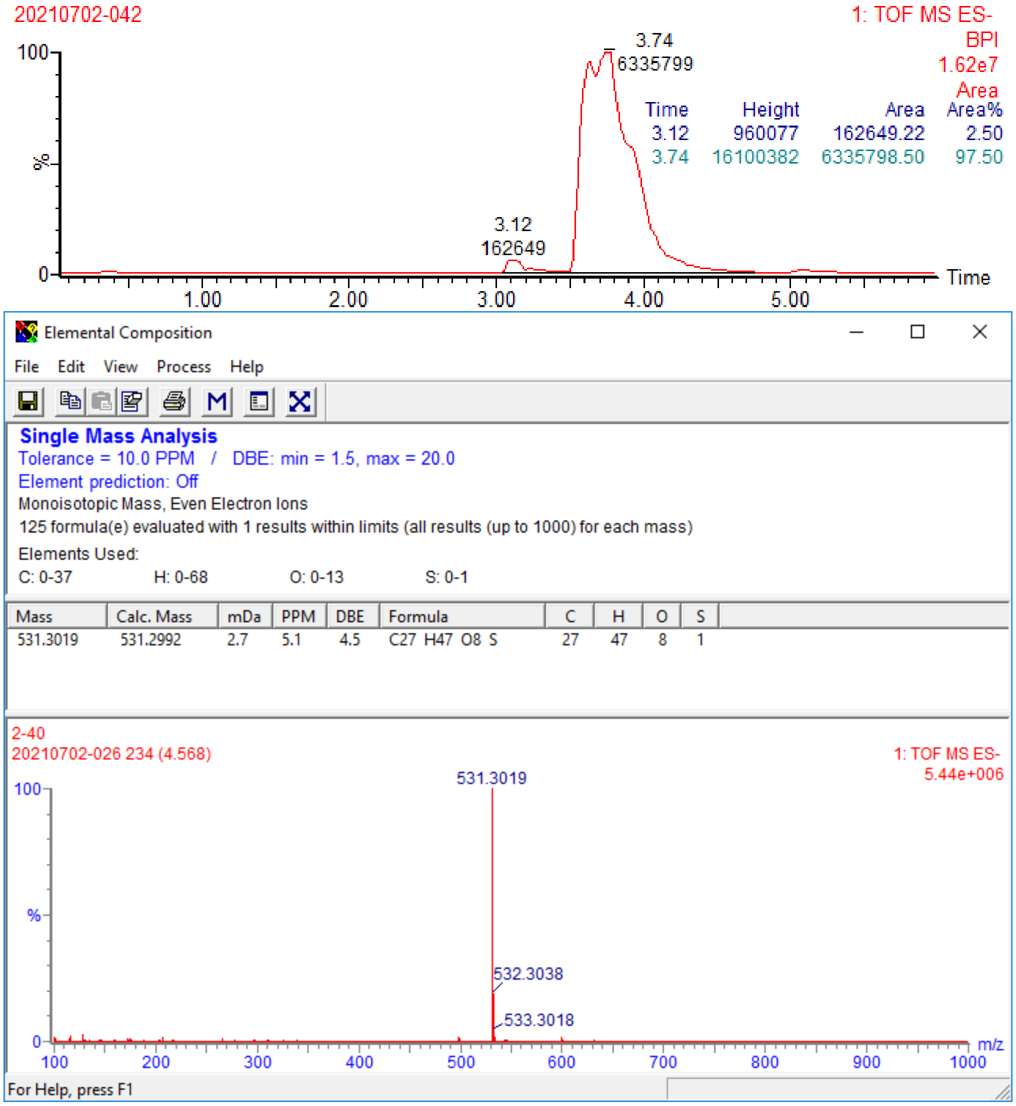

**Fig. S6** Purity and HR-ESI-MS of 5$\alpha$-cyprinol sulfate (**5CS**) on negative mode

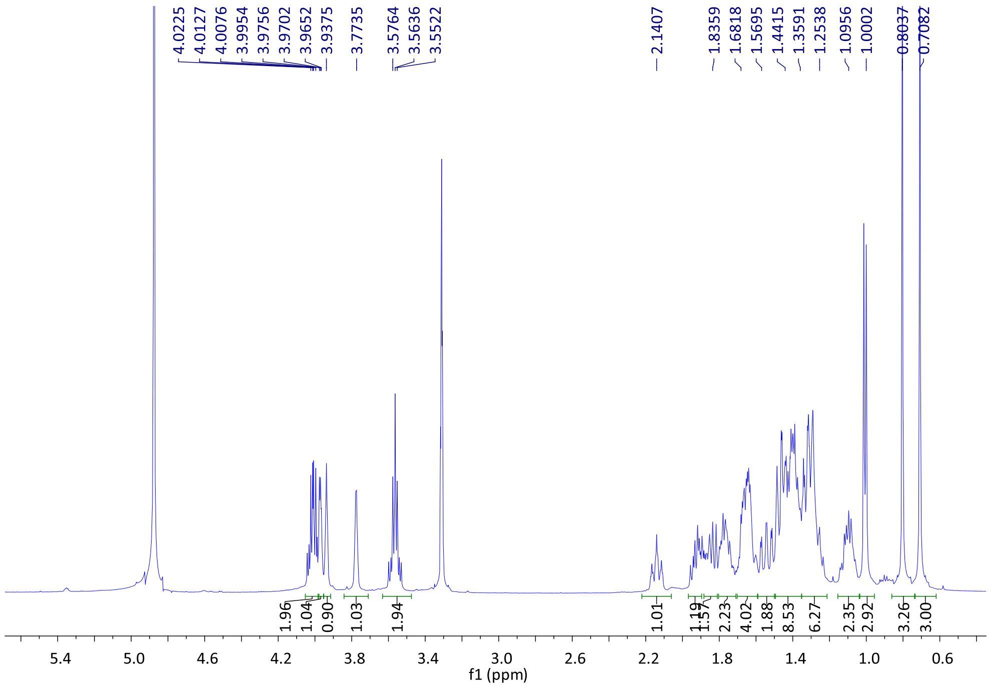


**Fig. S7** [1]H NMR spectrum of 5*α*-cyprinol sulfate **(5CS)**

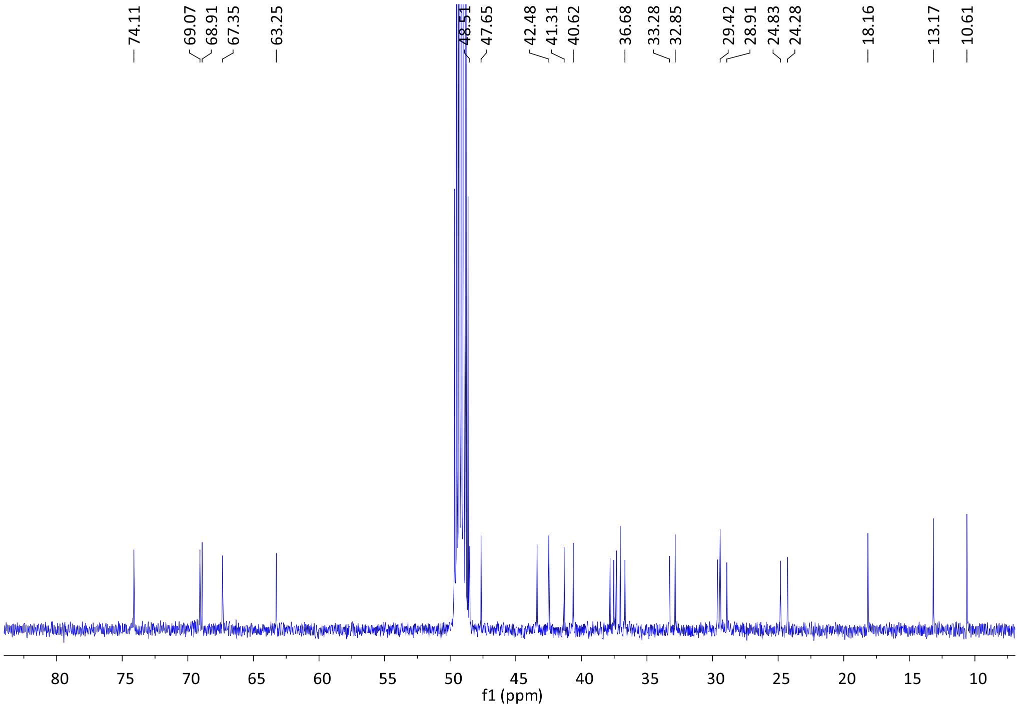


**Fig. S8** [13]C NMR spectrum of 5*α*-cyprinol sulfate **(5CS)**

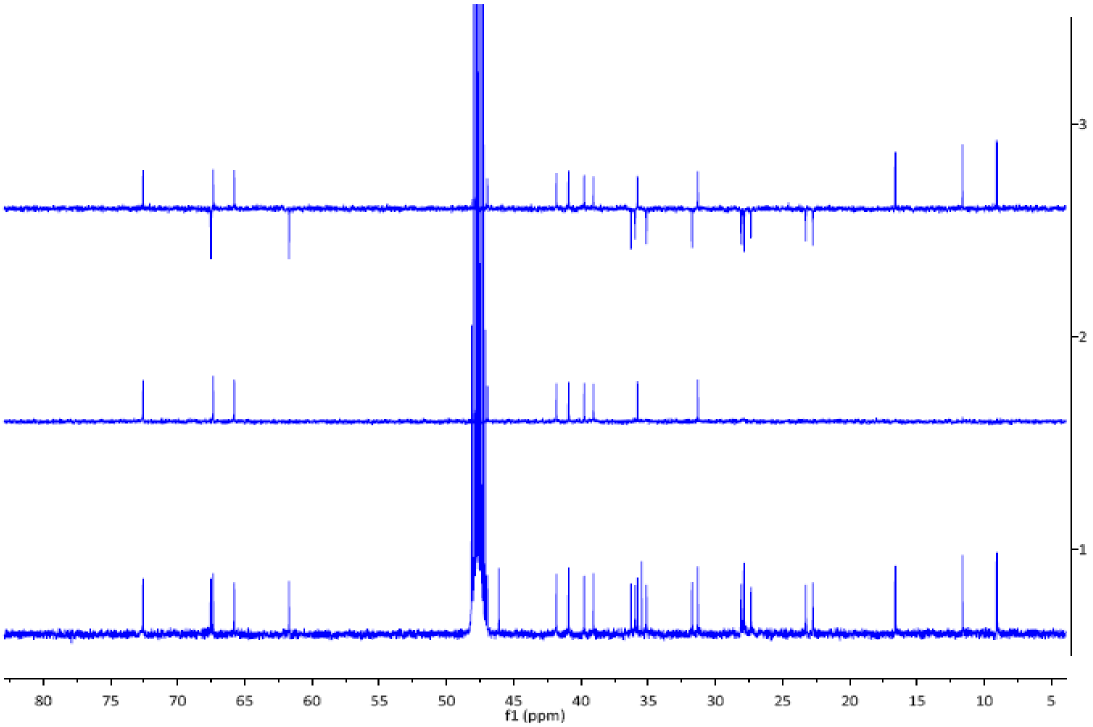


**Fig. S9** DEPT spectra of 5$\alpha$-cyprinol sulfate **(5CS)**

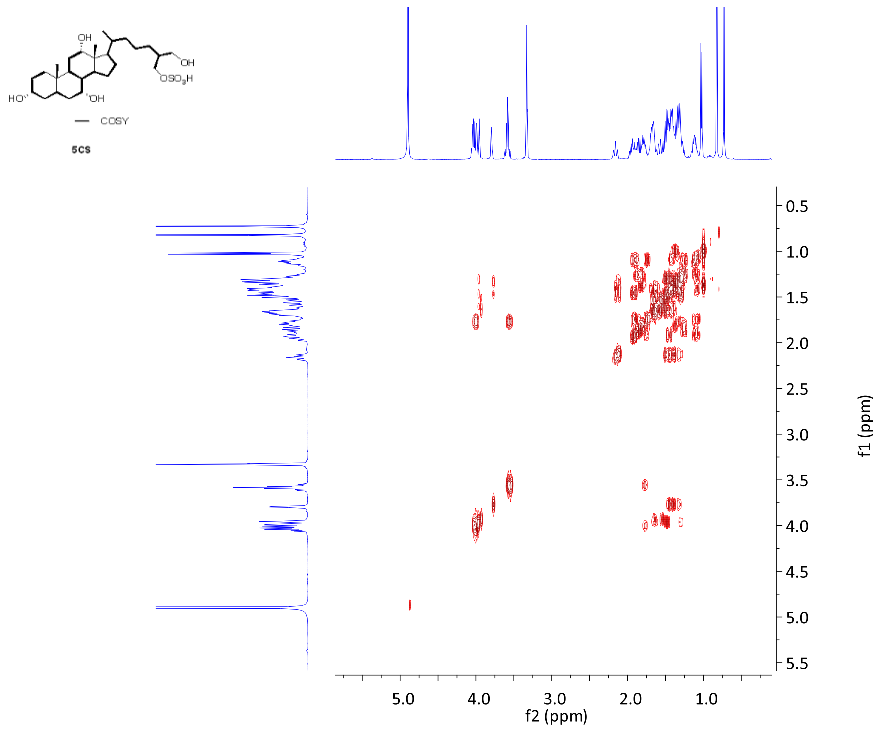


**Fig. S10** $^1H$-$^1H$ COSY spectrum of 5$\alpha$-cyprinol sulfate **(5CS)**

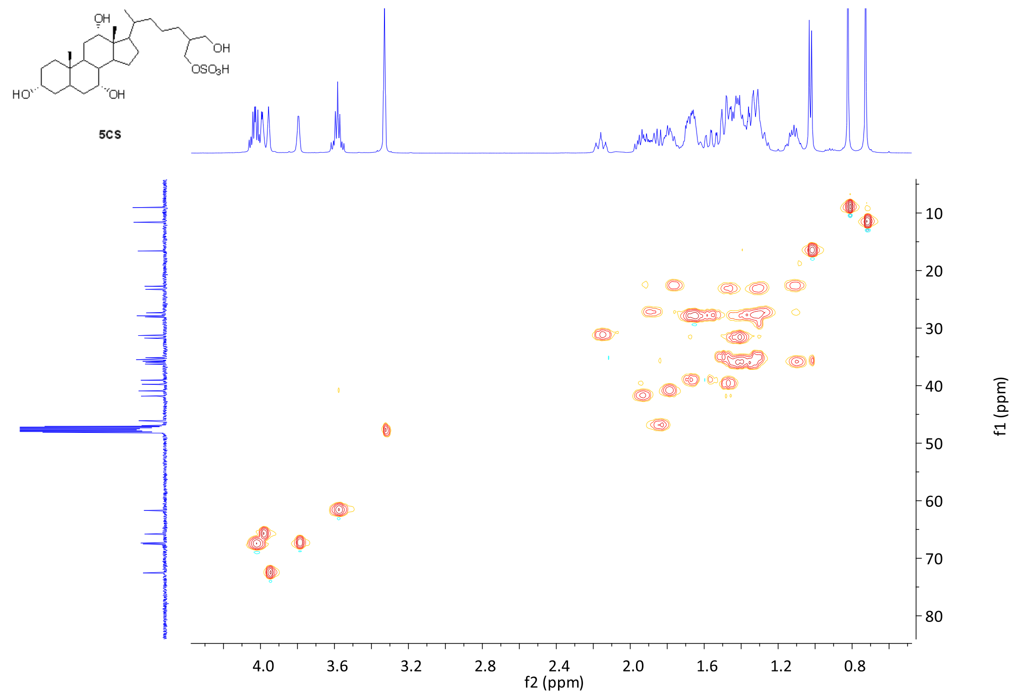


**Fig. S11** HSQC spectrum of 5$\alpha$-cyprinol sulfate **(5CS)**

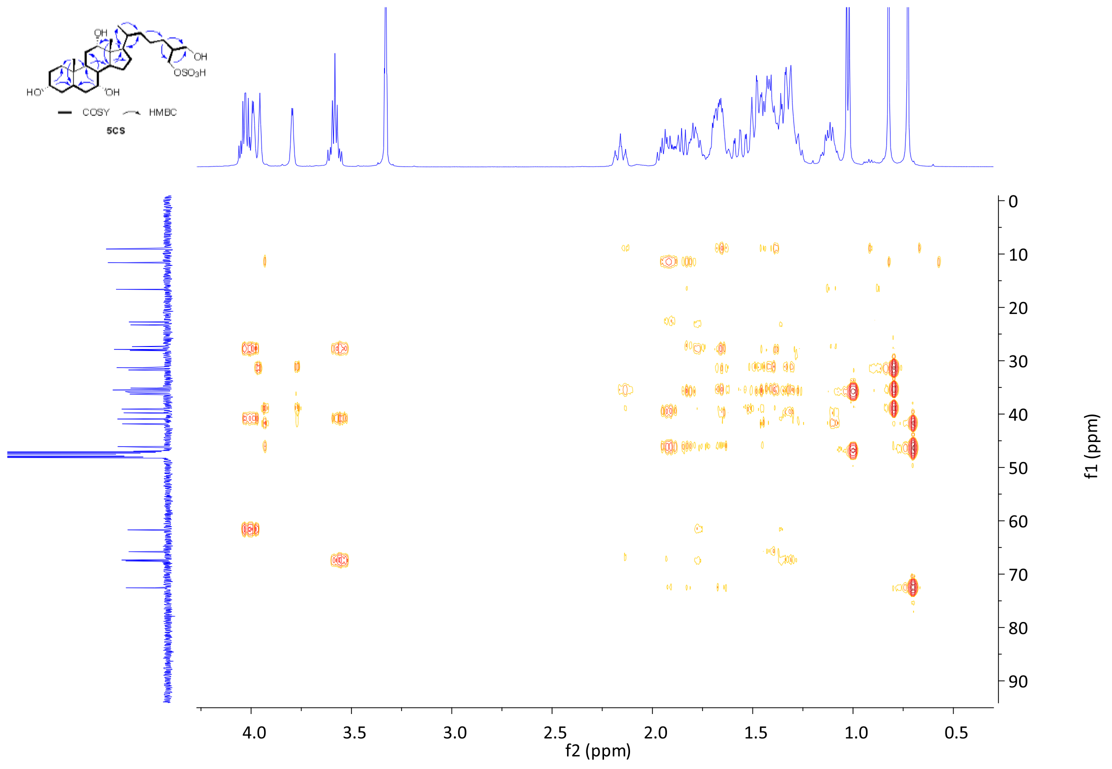


**Fig. S12** HMBC spectrum of 5$\alpha$-cyprinol sulfate **(5CS)**

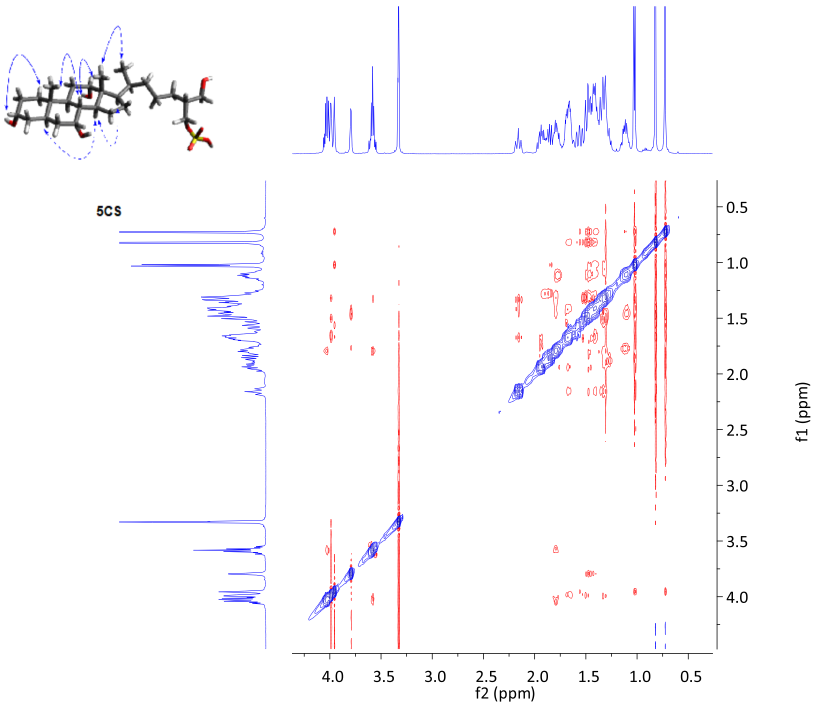


**Fig. S13** NOESY spectrum of 5$\alpha$-cyprinol sulfate **(5CS)**

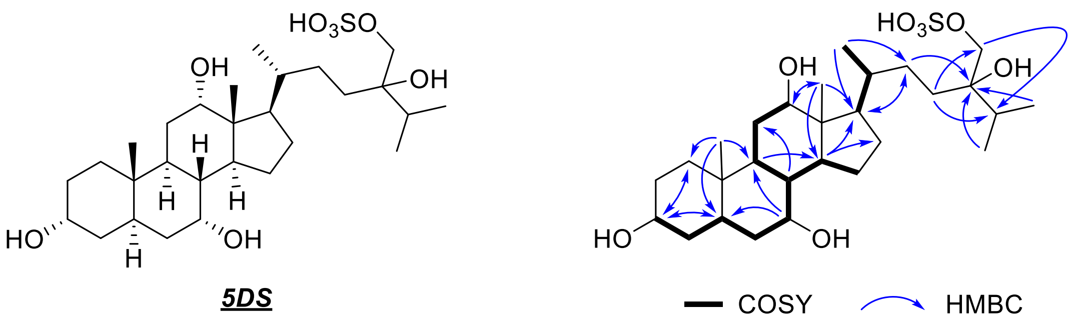


**Fig. S14** Key COSY and HMBC correlations of 5$\alpha$-daniol sulfate (**5DS**)

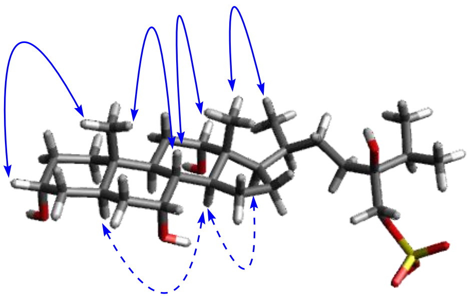

**Fig. S15** NOESY correlations of 5*α*-daniol sulfate (**5DS**)

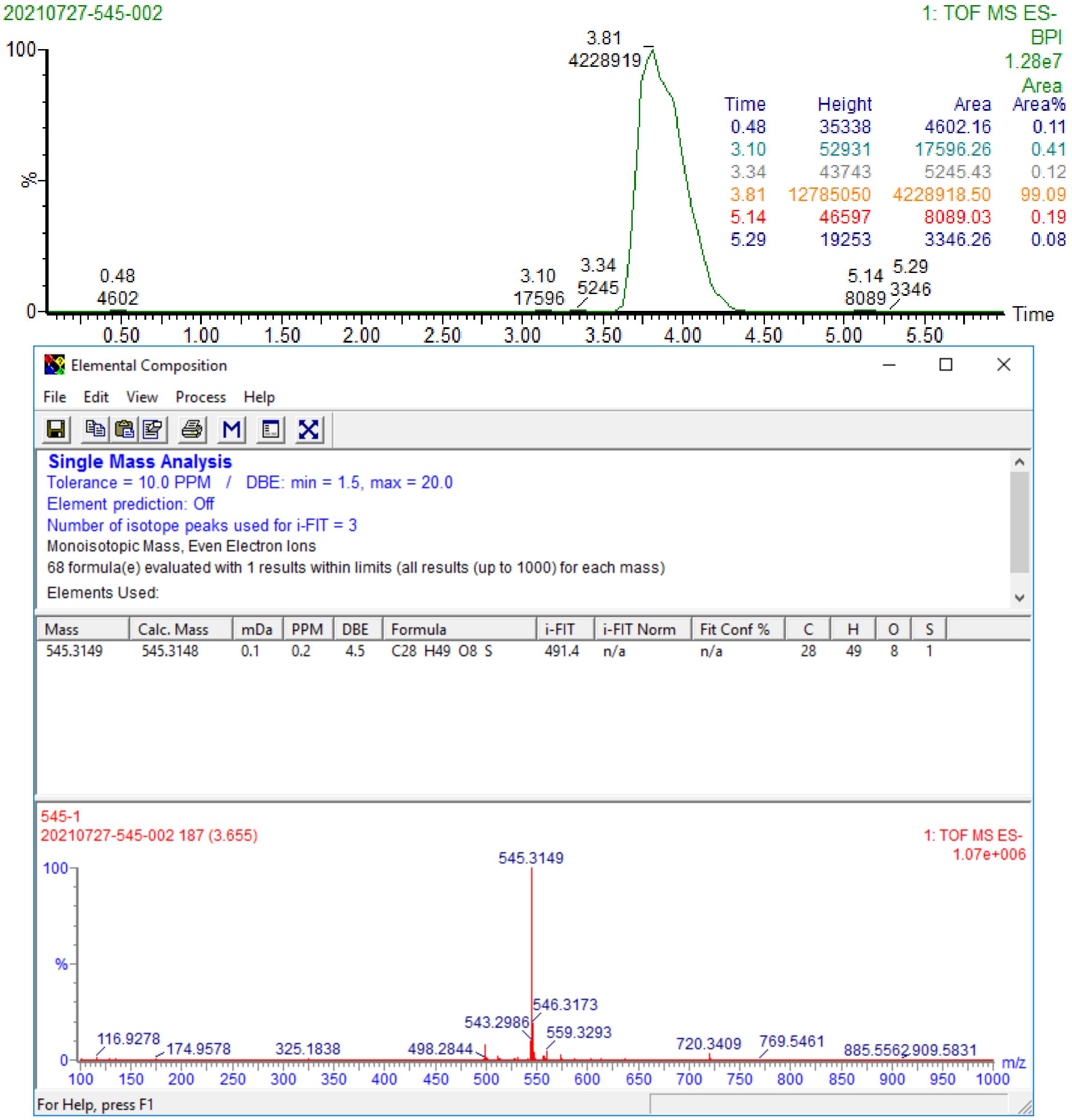


**Fig. S16** Purity and HR-ESI-MS of 5$\alpha$-daniol sulfate (**5DS**) on negative mode

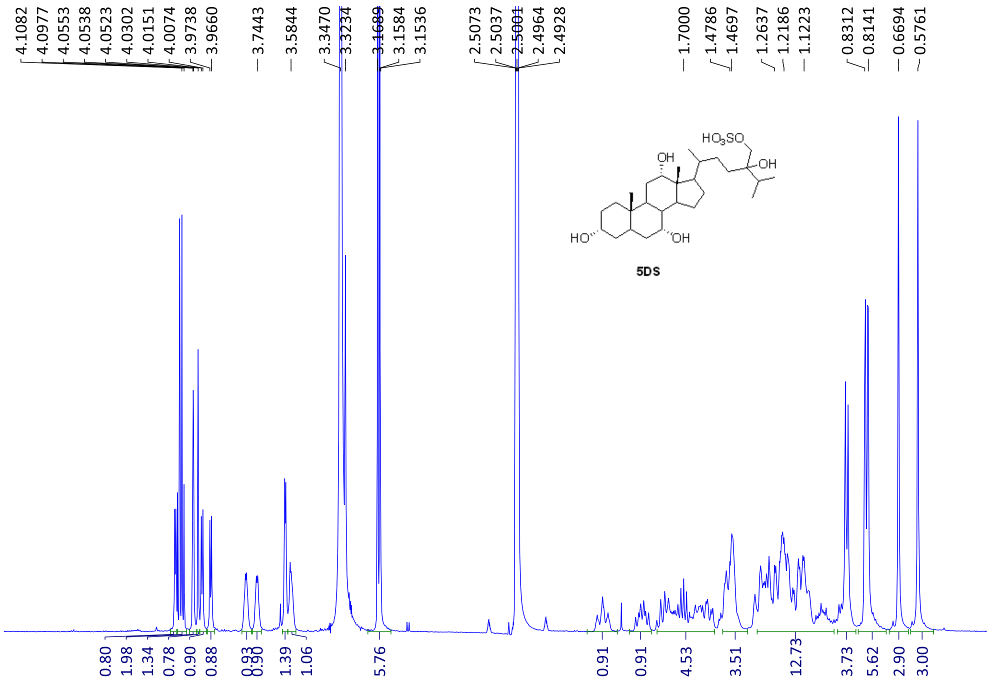

**Fig. S17** $^{1}$H NMR spectrum of 5$\alpha$-daniol sulfate (**5DS)**

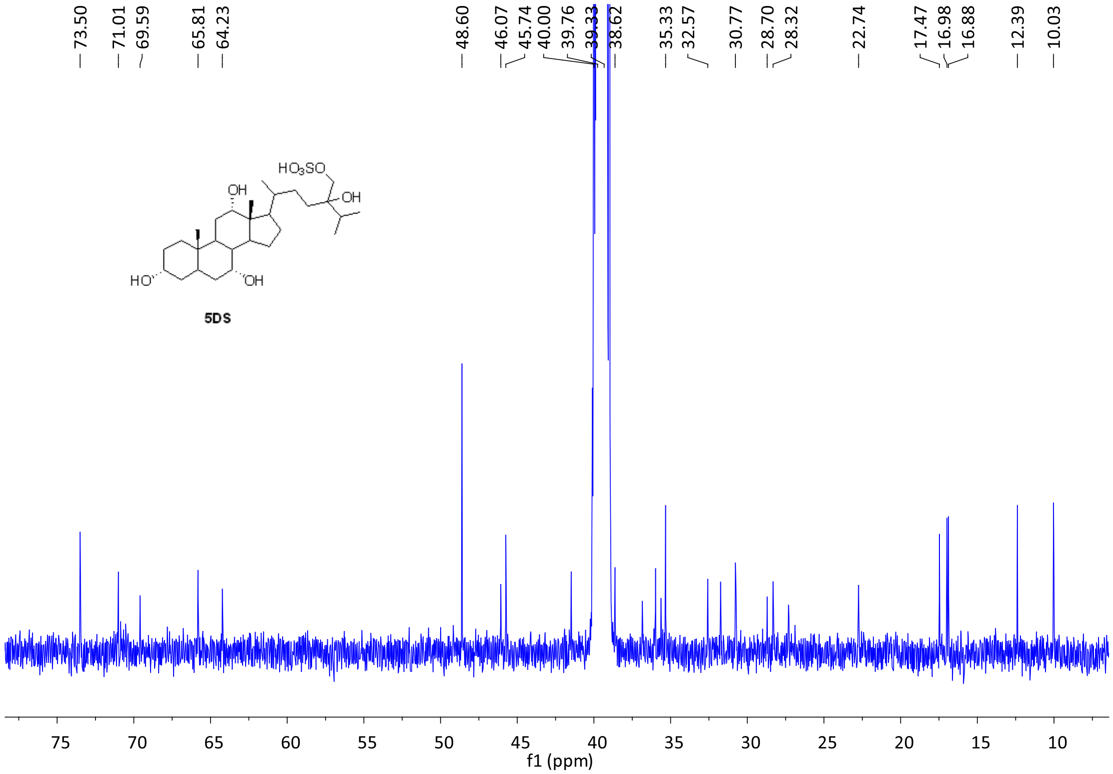


**Fig. S18** $^{13}$C NMR spectrum of 5$\alpha$-daniol sulfate (**5DS)**

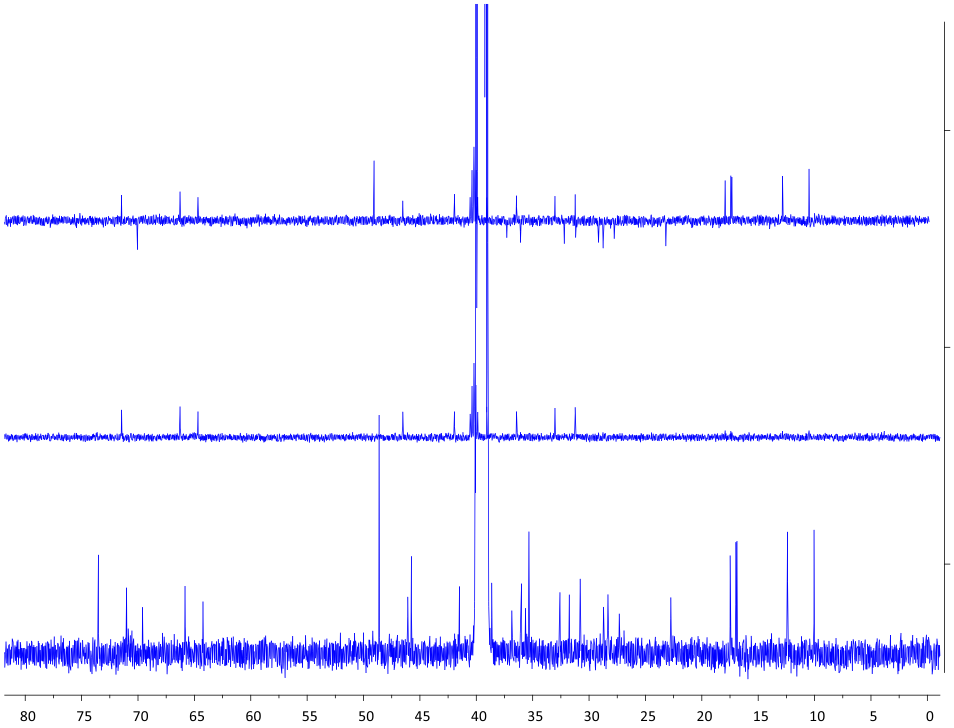


**Fig. S19** DEPT spectra of 5$\alpha$-daniol sulfate (**5DS**)

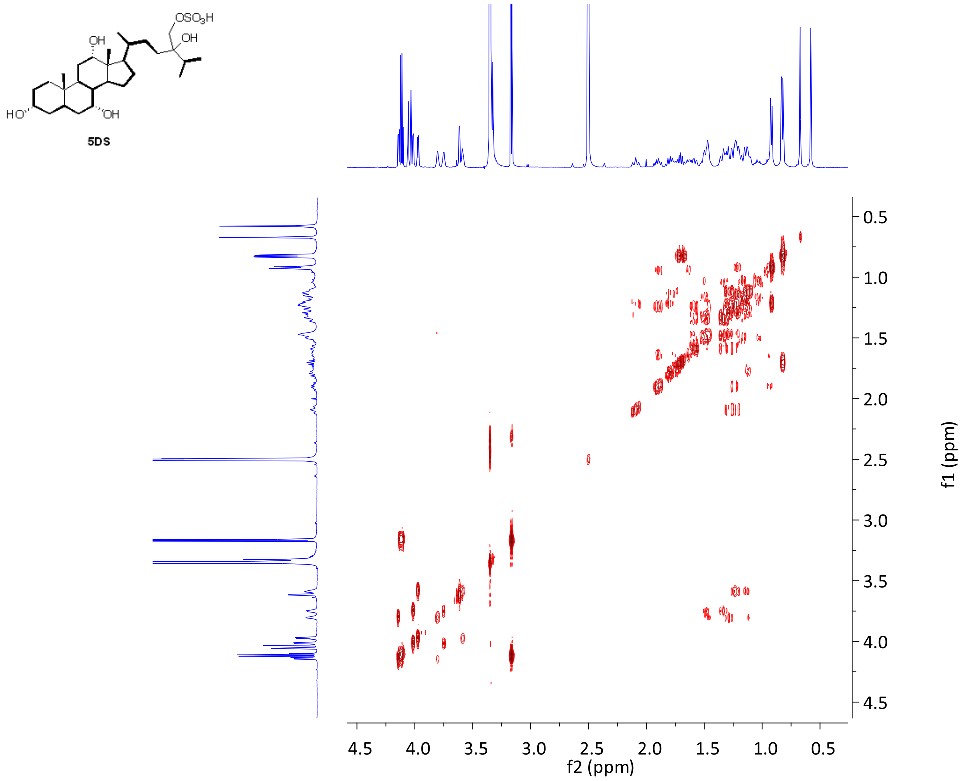


**Fig. S20** $^1$H-$^1$H COSY spectrum of 5$\alpha$-daniol sulfate (**5DS**)

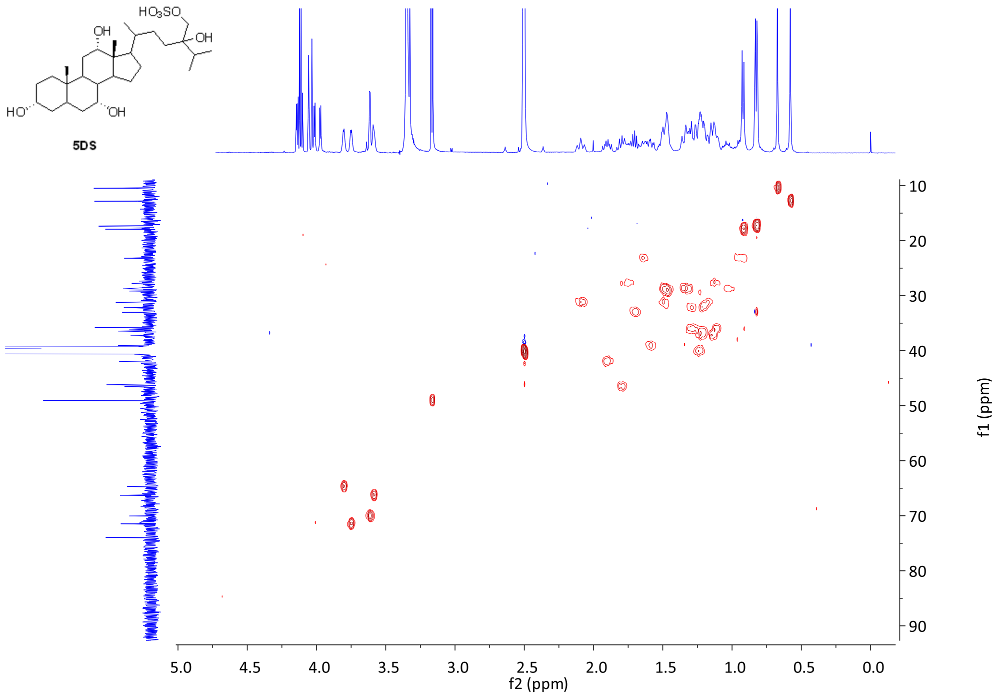


**Fig. S21** HSQC spectrum of 5*α*-daniol sulfate (**5DS)**

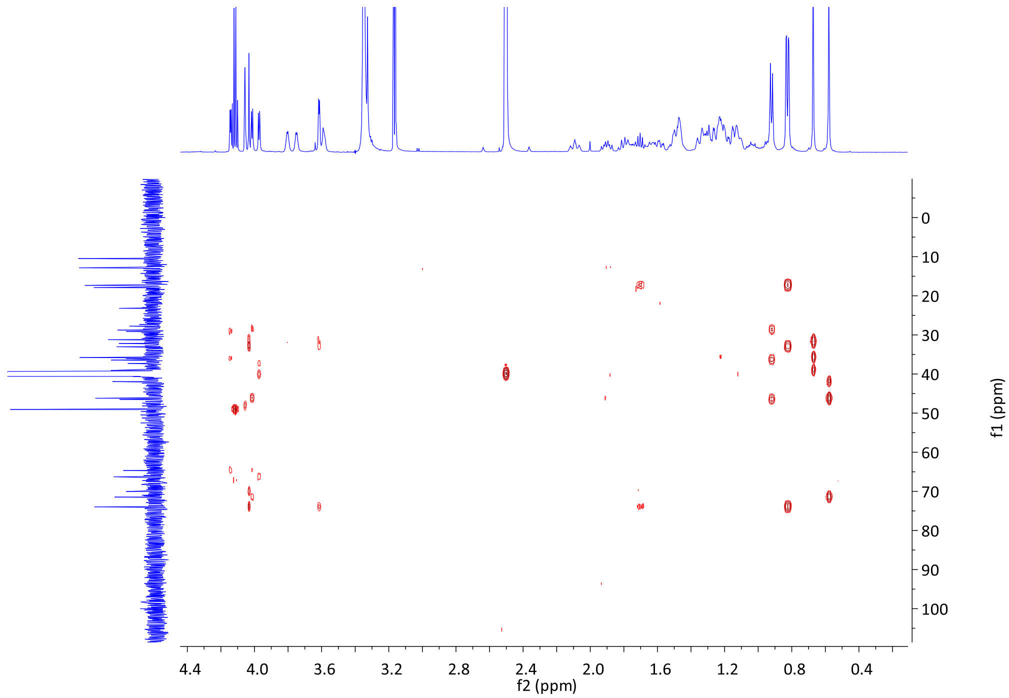


**Fig. S22** HMBC spectrum of 5*α*-daniol sulfate (**5DS)**

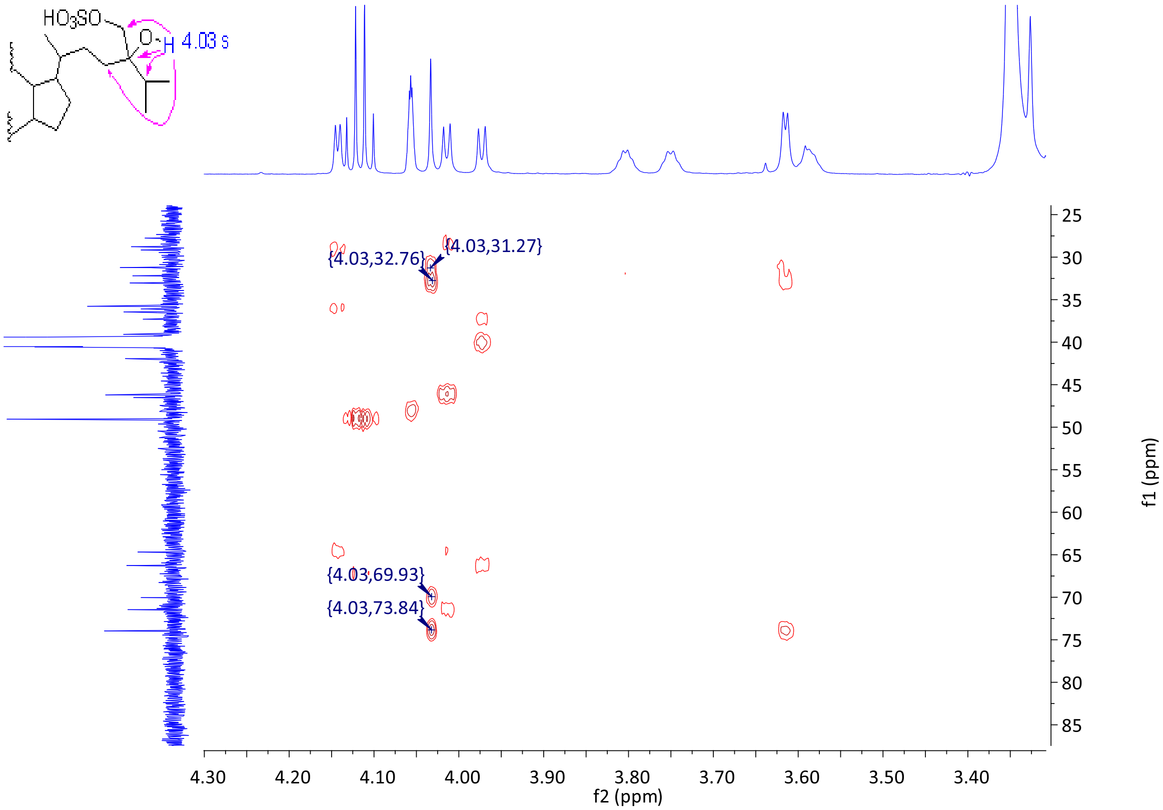


**Fig. S23** Partial zoom up of HMBC spectrum of 5$\alpha$-daniol sulfate (**5DS**)

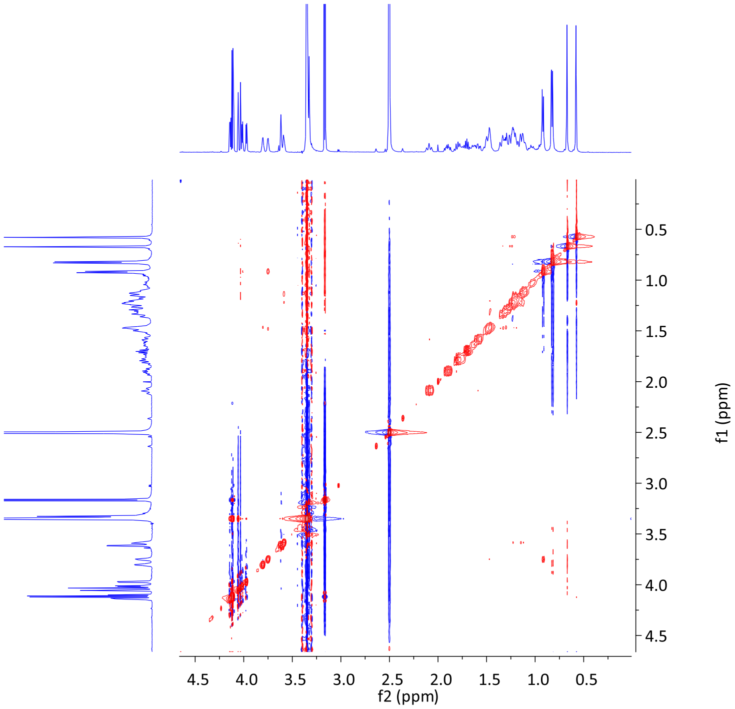


**Fig. S24** NOESY spectrum of 5$\alpha$-daniol sulfate (**5DS**)

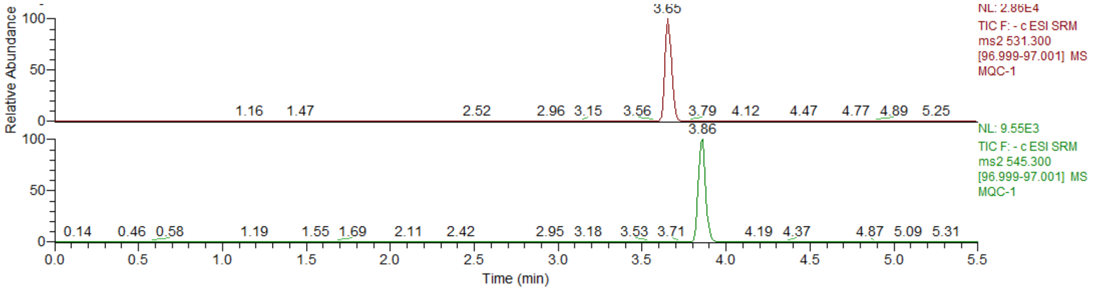


**Fig. S25** Chromatogram of oxysterol sulfates 5$\alpha$-cyprinol sulfate (**5CS**) and 5$\alpha$-daniol sulfate (**5DS**) in blank serum, concentration of each analyte is 8.0 ng/mL.

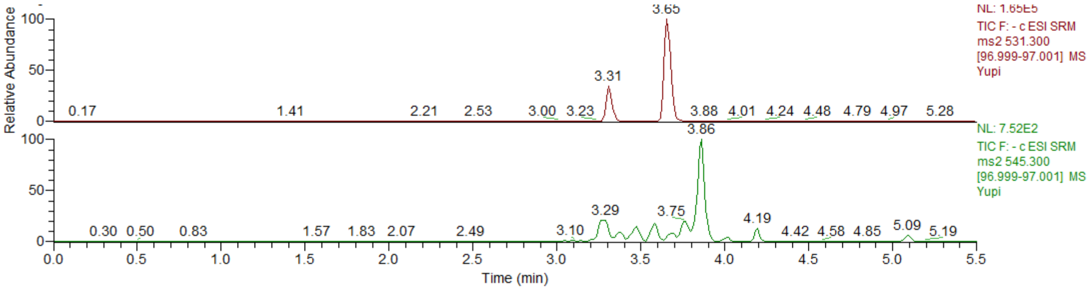


**Fig. S26** Chromatogram of oxysterol sulfates 5$\alpha$-cyprinol sulfate (**5CS**) and 5$\alpha$-daniol sulfate (**5DS)** in zebrafish skin extract (1000×SKE)

**Supplementary Table 1:** Statistical analysis of alarm behaviors in figure 1b-e (sig. = significant, n.s. = not significant, Con. = Control, Exp. = Experiment).

| **Erratic movements duration (%)** | | | | | | | | | | |
|---|---|---|---|---|---|---|---|---|---|---|
| Different phases | SKE | | nBuOH | | EtOAc | | PET | | Fraction 7 | |
| | Con. | Exp. | Con. | Exp. | Con. | Exp. | Con. | Exp. | Con. | Exp. |
| N of biologically independent samples | 9 | 8 | 14 | 14 | 15 | 15 | 15 | 15 | 8 | 8 |
| Tests of normality (sig. of Shapiro-Wilk) | 0.041 | 0.088 | 0.157 | 0.995 | 0.000 | 0.426 | 0.057 | 0.028 | 0.225 | 0.167 |
| Statistical analyses | Mann-Whitney U test | | Independent samples t-test | | Mann-Whitney U test | | Independent samples t-test | | | |
| *p* values (2-tailed) | $p<0.001$ | | $p<0.001$ | | n.s. | | n.s. | | $p<0.01$ | |

| **Time in bottom half (%)** | | | | | | | | | | |
|---|---|---|---|---|---|---|---|---|---|---|
| Different phases | SKE | | nBuOH | | EtOAc | | PET | | Fraction 7 | |
| | Con. | Exp. | Con. | Exp. | Con. | Exp. | Con. | Exp. | Con. | Exp. |
| N of biologically independent samples | 8 | 8 | 12 | 10 | 15 | 15 | 9 | 9 | 6 | 10 |
| Tests of normality (sig. of Shapiro-Wilk) | 0.007 | 0.969 | 0.145 | 0.443 | 0.079 | 0.254 | 0.297 | 0.185 | 0.674 | 0.267 |
| Statistical analyses | Mann-Whitney U test | | Independent samples t-test | | | | | | | |
| *p* values (2-tailed) | $p<0.01$ | | $p<0.01$ | | n.s. | | n.s. | | $p<0.001$ | |

| Freezing duration (%) | | | | | | | | | | |
|---|---|---|---|---|---|---|---|---|---|---|
| Different phases | SKE | | nBuOH | | EtOAc | | PET | | Fraction 7 | |
| | Con. | Exp. | Con. | Exp. | Con. | Exp. | Con. | Exp. | Con. | Exp. |
| N of biologically independent samples | 6 | 7 | 8 | 6 | 6 | 7 | 6 | 6 | 6 | 6 |
| Tests of normality (sig. of Shapiro-Wilk) | 0.727 | 0.096 | 0.000 | 0.891 | 0.093 | 0.050 | 0.545 | 0.645 | 0.243 | 0.939 |
| Statistical analyses *p* values (2-tailed) | Independent samples t-test | | Mann-Whitney U test | | Independent samples t-test | | | | | |
| | $p<0.001$ | | $p<0.01$ | | n.s. | | n.s. | | $p<0.001$ | |

| Latency to upper half (s) | | | | | | | | | | |
|---|---|---|---|---|---|---|---|---|---|---|
| Different phases | SKE | | nBuOH | | EtOAc | | PET | | Fraction 7 | |
| | Con. | Exp. | Con. | Exp. | Con. | Exp. | Con. | Exp. | Con. | Exp. |
| N of biologically independent samples | 6 | 6 | 6 | 6 | 12 | 9 | 6 | 6 | 6 | 6 |
| Tests of normality (sig. of Shapiro-Wilk) | 0.533 | 0.008 | 0.869 | 0.366 | 0.458 | 0.671 | 0.488 | 0.651 | 0.890 | 0.876 |
| Statistical analyses *p* values (2-tailed) | Mann-Whitney U test | | Independent samples t-test | | | | | | | |
| | $p<0.01$ | | $p<0.001$ | | n.s. | | n.s. | | $p<0.001$ | |

**Supplementary Table 2:**[1]H NMR assignments ($\delta$ in ppm, $J$ in hertz present in parentheses) for isolated compound 5$\alpha$-cyprinol sulfate (**5CS**) in method-$d_4$, 5$\alpha$-cyprinol sulfate [8] in method-$d_4$, 5$\beta$-cyprinol sulfate [9] in $C_5H_5N$-$d_5$, and petromyzonol sulfate [2] in method-$d_4$

| No. | **5CS** | 5$\alpha$-cyprinol sulfate | 5$\beta$-cyprinol sulfate | petromyzonol sulfate |
|---|---|---|---|---|
| 1$\beta$ | 1.40, m | 1.42, m | N.A | 1.39, m |
| 1$\alpha$ | 1.40, m | 1.42, m | N.A | 1.43, m |
| 2$\beta$ | 1.65, m[a] | 1.64, m[a] | N.A | 1.62, m[a] |
| 2$\alpha$ | 1.64, m[a] | 1.64, m[a] | N.A | 1.66, m[a] |
| 3$\beta$ | 3.97, m | 3.99, dddd (2.6) | 3.35, m | 3.97, m |
| 4$\beta$ | 1.32, m | 1.32, m | N.A | 1.31, m |
| 4$\alpha$ | 1.49, m | 1.51, m | N.A | 1.49, ddd (14.0, 14.0, 3.0) |
| 5$\alpha$ | 2.14, ddd (13.0, 6.1, 3.0) | 2.15, tt (13.0, 3.2) | N.A | 2.14, dddd (13.0, 13.0, 3.0, 3.0) |
| 6$\beta$ | 1.35, m | 1.35, m | N.A | 1.42, m |
| 6$\alpha$ | 1.42, m | 1.42, m | N.A | 1.32, ddd (14.0, 3.0, 3.0) |
| 7$\beta$ | 3.78, m | 3.79, dd (2.7) | 3.78, m | 3.78, ddd (3.0, 3.0, 3.0) |
| 8$\beta$ | 1.46, m | 1.48, m | N.A | 1.46, ddd (11.5, 11.5, 3.0) |
| 9$\alpha$ | 1.66, m | 1.68, m | N.A | 1.66, m |
| 11$\beta$ | 1.56, m | 1.57, m | N.A | 1.55, m |
| 11$\alpha$ | 1.65, m | 1.68, m | N.A | 1.66, m |
| 12$\beta$ | 3.94, m | 3.95, dd (2.6) | 3.95, m | 3.93, dd (3.0, 3.0) |
| 14$\alpha$ | 1.92, m | 1.94, dt (12.1, 7.3) | N.A | 1.93, ddd (12.0, 12.0, 7.5) |
| 15$\beta$ | 1.11, m | 1.12, m | N.A | 1.11, dddd (12.0, 12.0, 12.0, 6.0) |
| 15$\alpha$ | 1.75, m | 1.77, m | N.A | 1.75, m |
| 16$\beta$ | 1.30, m | 1.28, m | N.A | 1.89, m |
| 16$\alpha$ | 1.75, m | 1.89, m | N.A | 1.28, m |
| 17$\alpha$ | 1.81, m | 1.85, t (9.3) | N.A | 1.84, ddd (9.5, 9.5, 9.5) |
| 18 | 0.71, m | 0.72, s | 0.71, s | 0.71, s |
| 19 | 0.80, m | 0.82, s | 0.91, s | 0.80, s |
| 20 | 1.39, m | 1.40, m | N.A | 1.41, m |
| 21 | 1.01, d (6.5) | 1.02, d (6.6) | 1.01, d (6.5) | 1.02, d (6.5) |
| 22a | 1.11, m | 1.11, m | N.A | 1.15, m |
| 22b | 1.45, m | 1.46, m | N.A | 1.54, m |
| 23a | 1.30, m | 1.31, m | N.A | 1.56, m |
| 23b | 1.45, m | 1.48, m | N.A | 1.75, m |
| 24a | 1.31, m | 1.35, m | N.A | 3.97, m |
| 24b | 1.31, m | 1.35, m | N.A | 3.97, m |
| 25 | 1.77, m | 1.80, m | N.A | absent |
| 26a | 4.01, m | 4.05, ddd (9.8, 6.1, 4.7) [a] | 3.55, d (6.0) [a] | absent |

| | | | | |
|---|---|---|---|---|
| 26b | 4.01, m | 4.00, ddd (9.8, 6.1, 4.7) [a] | 3.55, d (6.0) [a] | absent |
| 27a | 3.56, m | 3.60, ddd (11.1, 6.1, 5.3) | 3.55, d (6.0) | absent |
| 27b | 3.56, m | 3.56, ddd (11.1, 6.1, 5.3) | 3.55, d (6.0) | absent |

[a] Resonances assigned to $H_{\alpha}$ and $H_{\beta}$ may be interchanged. N.A, not assigned in reference.

**Supplementary Table 3:** $^{13}$C NMR assignments (*δ* in ppm) for isolated compound 5α-cyprinol sulfate (**5CS**) in method-$d_4$, 5*α*-cyprinol sulfate [8] in method-$d_4$, 5*β*-cyprinol sulfate [9] in $C_5H_5N$-$d_5$, and petromyzonol sulfate [2] in method-$d_4$

| No. | **5CS** | 5*α*-cyprinol sulfate | 5*β*-cyprinol sulfate | petromyzonol sulfate |
|---|---|---|---|---|
| 1 | 33.3 ($CH_2$) | 31.7 ($CH_2$) | 37.3 ($CH_2$) | 32.9 ($CH_2$) |
| 2 | 28.1 ($CH_2$) | 28.1 ($CH_2$) | 32.2 ($CH_2$) | 29.2 ($CH_2$) |
| 3 | 67.4 (CH) | 65.3 (CH) | 73.7 (CH) | 66.9 (CH) |
| 4 | 36.7 ($CH_2$) | 35.1 ($CH_2$) | 41.3 ($CH_2$) | 36.3 ($CH_2$) |
| 5 | 32.9 (CH) | 31.2 (CH) | 43.8 (CH) | 32.5 (CH) |
| 6 | 37.5 ($CH_2$) | 36.3 ($CH_2$) | 36.7 ($CH_2$) | 37.4 ($CH_2$) |
| 7 | 68.9 (CH) | 67.4 (CH) | 69.9 (CH) | 68.5 (CH) |
| 8 | 41.3 (CH) | 39.8 (CH) | 41.8 (CH) | 40.9 (CH) |
| 9 | 40.6 (CH) | 39.0 (CH) | 28.7 (CH) | 40.2 (CH) |
| 10 | 37.8 (qC) | 35.5 (qC) | 36.7 (qC) | 36.6 (qC) |
| 11 | 29.6[a] ($CH_2$) | 27.9 ($CH_2$) | 29.6 ($CH_2$) | 29.2 ($CH_2$) |
| 12 | 74.1 (CH) | 72.6 (CH) | 74.9 (CH) | 73.7 (CH) |
| 13 | 47.7 (qC) | 46.0 (qC) | 48.2 (qC) | 47.1 (qC) |
| 14 | 43.4 (CH) | 41.7 (CH) | 44.0 (CH) | 42.9 (CH) |
| 15 | 24.3 ($CH_2$) | 22.7 ($CH_2$) | 25.0 ($CH_2$) | 23.9 ($CH_2$) |
| 16 | 29.4[a] ($CH_2$) | 27.3 ($CH_2$) | 30.4 ($CH_2$) | 28.4 ($CH_2$) |
| 17 | 48.5 (CH) | 47.1 (CH) | 49.1 (CH) | 48.0 ($CH_2$) |
| 18 | 13.2 ($CH_3$) | 11.5 ($CH_3$) | 13.8 ($CH_3$) | 12.7 ($CH_3$) |
| 19 | 10.6 ($CH_3$) | 8.9 ($CH_3$) | 24.0 ($CH_3$) | 10.2 ($CH_3$) |
| 20 | 37.0 (CH) | 35.8 (CH) | 38.0 (CH) | 36.6 (CH) |
| 21 | 18.2 ($CH_3$) | 16.7 ($CH_3$) | 18.9 ($CH_3$) | 17.6 ($CH_3$) |
| 22 | 37.3 ($CH_2$) | 36.0 ($CH_2$) | 38.3 ($CH_2$) | 32.8 ($CH_2$) |
| 23 | 24.8 ($CH_2$) | 23.3 ($CH_2$) | 25.6 ($CH_2$) | 27.0 ($CH_2$) |
| 24 | 29.4[a] ($CH_2$) | 27.8 ($CH_2$) | 30.2 ($CH_2$) | 69.4 ($CH_2$) |
| 25 | 42.5 (CH) | 40.9 (CH) | 45.2 (CH) | absent |
| 26 | 69.1 ($CH_2$) | 67.6 ($CH_2$) | 68.5 ($CH_2$) | absent |
| 27 | 63.3 ($CH_2$) | 61.7 ($CH_2$) | 64.6 ($CH_2$) | absent |

[a] chemical shifts are interchangeable

**Supplementary Table 4:** $^{1}H$ and $^{13}C$ NMR assignments ($\delta$ in ppm, $J$ in hertz present in parentheses) for isolated compound **5DS** in DMSO-$d_6$.

| No. | $\delta_H$ ($J$ in Hz) | $\delta_C$ | No. | $\delta_H$ ($J$ in Hz) | $\delta_C$ |
|---|---|---|---|---|---|
| 1 | 1.48, 1.18 (m) | 31.2 ($CH_2$) | 17 | 1.82 (m) | 46.1 (CH) |
| 2 | 1.47, 1.33 (m) | 28.7 ($CH_2$) | 18 | 0.58 (s) | 12.4 ($CH_3$) |
| 3 | 3.79 (m) | 64.2 (CH) | 19 | 0.67 (s) | 10.0 ($CH_3$) |
| 4 | 1.28, 1.12 (m) | 35.6 ($CH_2$) | 20 | 1.28 (m) | 36.0 (CH) |
| 5 | 2.08 (m) | 30.7 (CH) | 21 | 0.92 (d, 6.5) | 17.5 ($CH_3$) |
| 6 | 1.28, 1.12 (m) | 36.8 ($CH_2$) | 22 | 1.48, 1.33 (m) | 28.3 ($CH_2$) |
| 7 | 3.58 (m) | 65.8 (CH) | 23 | 1.49, 1.31 (m) | 30.8 ($CH_2$) |
| 8 | 1.59 (m) | 39.5 (CH) | 24 | --- | 73.5 (qC) |
| 9 | 1.22 (m) | 38.9 (CH) | 25 | 1.70 (m) | 32.6 (CH) |
| 10 | --- | 35.3 (qC) | 26 | 0.82 (d, 6.9), | 17.0 ($CH_3$) |
| 11 | 1.73, 1.14 (m) | 28.3 ($CH_2$) | 27 | 0.83 (d, 6.9) | 16.9 ($CH_3$) |
| 12 | 3.75 (m) | 71.0 (CH) | 28 | 3.62 (d, 2.5) | 69.6 ($CH_2$) |
| 13 | --- | 45.7 (qC) | OH-3 | 4.14 (d, 2.8) | |
| 14 | 1.90 (m) | 41.5 (CH) | OH-7 | 3.97 (d, 3.9) | |
| 15 | 1.64, 0.97 (m) | 22.7 ($CH_2$) | OH-12 | 4.01 (d, 3.9) | |
| 16 | 1.47, 1.34 (m) | 28.7 ($CH_2$) | OH-24 | 4.03 (s) | |

**Supplementary Table 5:** MS parameters for 5$\alpha$-cyprinol sulfate (**5CS**) and 5$\alpha$-daniol sulfate (**5DS**) on negative mode

| Analytes | Retention time (min) | Molecular weight | Molecular formula | Q1 (Da) | Q3 (Da) | Collision energy (eV) | RF-Lens （V） |
|---|---|---|---|---|---|---|---|
| **5CS** | 3.65 | 532.73 | $C_{27}H_{46}O_8S$ | 531.3 | 97.0 | 55 | 190 |
| **5DS** | 3.86 | 546.73 | $C_{27}H_{48}O_7S$ | 545.3 | 97.0 | 55 | 190 |

**Supplementary Table 6:** Linearity, LLOQ, and LOD for 5$\alpha$-cyprinol sulfate (**5CS**) and 5$\alpha$-daniol sulfate (**5DS**)

| Analytes | Linear range (ng/mL) | Linear equation | $r^2$ | LLOQ (ng/mL) | LOD (ng/mL) |
|---|---|---|---|---|---|
| **5CS** | 0.2-100 | $y = 10574\chi - 5656$ | 0.9992 | 0.20 | 0.12 |
| **5DS** | 0.2-100 | $y=3700.1\chi - 1242.3$ | 0.9996 | 0.20 | 0.12 |

**Supplementary Table 7:** Intraday and interday accuracy and precision of the LC-MS/MS method for 5$\alpha$-cyprinol sulfate (**5CS**) and 5$\alpha$-daniol sulfate (**5DS**)

| Analyte | QC level | Calc conc. (mean ± SD ng/mL) | | Accuracy (DEV, %) | | Precision (RSD, %) | |
|---|---|---|---|---|---|---|---|
| | | Intra-day | Inter-day | Intra-day | Inter-day | Intra-day | Inter-day |
| **5CS** | LQC | 0.62±0.01 | 0.59±0.03 | 102.07% | 98.91% | 1.18 | 4.32 |
| | MQC | 8.08±0.25 | 7.82±0.24 | 101.02% | 97.82% | 3.11 | 3.02 |
| | HQC | 84.02±1.77 | 86.4±2.65 | 105.03% | 108.03% | 2.11 | 3.07 |
| **5DS** | LQC | 0.62±0.02 | 0.59±0.02 | 103.05% | 98.57% | 3.40 | 3.94 |
| | MQC | 8.05±0.33 | 7.80±0.21 | 100.60% | 97.59% | 4.15 | 2.71 |
| | HQC | 82.31±1.91 | 84.18±2.11 | 102.89% | 105.23% | 2.32 | 2.51 |

**Note**: nominal concentration for LQC, MQC, and HQC were designed as 0.6 ng/mL, 8.0 ng/mL, and 80.0 ng/mL, respectively.

**Supplementary Table 8:** The concentration of 5$\alpha$-cyprinol sulfate (**5CS**) and 5$\alpha$-daniol sulfate (**5DS**) in a 1000×SKE stock solution was determined using the LC-MS/MS method. The stock solution of 1000×SKE was diluted by a factor of 100 in order to fall within the linear range of the analytical method. The actual concentration should be multiplied by a factor of 100.

| Analyte | | Test 1 | Test 2 | Test 3 | Test 4 | Test 5 | Average | SD |
|---|---|---|---|---|---|---|---|---|
| **5CS** | (ng/mL) | 535.63 | 542.40 | 545.40 | 527.50 | 527.20 | 535.63 | 8.34 |
| | (nmol/mL) | 1.0068 | 1.0195 | 1.0252 | 0.9915 | 0.9910 | 1.0100 | 0.02 |
| **5DS** | (ng/mL) | 5.36 | 5.56 | 5.67 | 5.46 | 5.67 | 5.54 | 0.14 |
| | (nmol/mL) | $0.98\times10^{-2}$ | $1.02\times10^{-2}$ | $1.04\times10^{-2}$ | $1.00\times10^{-2}$ | $1.04\times10^{-2}$ | $1.02\times10^{-2}$ | $0.025\times10^{-2}$ |

**Supplementary Table 9:** Statistical analysis of alarm behaviors in figure 3a-h (n.s. = not significant).

**Erratic movements duration (%) caused by 5CS (5DS)**
**Bonferroni test**
**5CS: ANOVA F (4, 51) = 11.891, $p<0.001$; 5DS: ANOVA F (4, 54) = 12.618, $p<0.001$**
**5CS Error: Between MS = 5.786, df =4; 5DS Error: Between MS = 1.230, df =4**

| | Fraction | 0 | –12 | –10 | –8 | –6 |
|---|---|---|---|---|---|---|
| N of biologically independent samples | | 13<br>(13) | 8<br>(11) | 9<br>(12) | 10<br>(9) | 16<br>(14) |
| | 0 | | n.s.<br>($p<0.01$) | $p<0.05$<br>(n.s.) | $p<0.001$<br>(n.s.) | n.s.<br>($p<0.001$) |
| | –12 | n.s.<br>($p<0.01$) | | n.s.<br>($p<0.001$) | $p<0.001$<br>($p<0.001$) | n.s.<br>(n.s.) |
| | –10 | $p<0.05$<br>(n.s.) | n.s.<br>($p<0.001$) | | $p<0.05$<br>(n.s.) | n.s.<br>($p<0.001$) |
| | –8 | $p<0.001$<br>(n.s.) | $p<0.001$<br>($p<0.001$) | $p<0.05$<br>(n.s.) | | $p<0.001$<br>($p<0.001$) |
| | –6 | n.s.<br>($p<0.001$) | n.s.<br>(n.s.) | n.s.<br>($p<0.001$) | $p<0.001$<br>($p<0.001$) | |

**Erratic movements duration (%) caused by 5CS+5DS**
**Tamhane test**
**Significance of homogeneity test of variance: $p<0.05$**

| Fraction | 0 | –12 | –11 | –10 | –9 | SKE |
|---|---|---|---|---|---|---|
| | 13 | 7 | 10 | 8 | 10 | 8 |
| 0 | | $p<0.05$ | $p<0.001$ | $p<0.05$ | $p<0.001$ | $p<0.001$ |
| –12 | $p<0.05$ | | n.s. | n.s. | $p<0.05$ | $p<0.01$ |
| –11 | $p<0.001$ | n.s. | | $p<0.01$ | $p<0.05$ | $p<0.01$ |
| –10 | $p<0.05$ | n.s. | $p<0.01$ | | $p<0.001$ | $p<0.001$ |
| –9 | $p<0.001$ | $p<0.05$ | $p<0.05$ | $p<0.001$ | | n.s. |
| SKE | $p<0.001$ | $p<0.01$ | $p<0.01$ | $p<0.001$ | n.s. | |

**Time in bottom half (%) caused by 5CS (5DS)**
**Kruskal-Wallis test of 5CS: H (4, N= 88)=29.419, $p$ <0.001;**
**Kruskal-Wallis test of 5DS: H (4, N= 88)=23.844, $p$ <0.001**
**Multiple Comparisons $p$ values (2-tailed) with Bonferroni correction**

| | Fraction | 0 | –12 | –10 | –8 | –6 |
|---|---|---|---|---|---|---|
| N of biologically independent samples | | 8<br>(8) | 20<br>(20) | 20<br>(20) | 20<br>(20) | 20<br>(20) |
| | 0 | | $p$<0.01<br>($p$<0.01) | $p$<0.001<br>($p$<0.05) | $p$<0.001<br>($p$<0.01) | n.s.<br>($p$<0.001) |
| | –12 | $p$<0.01<br>($p$<0.01) | | n.s.<br>(n.s.) | n.s.<br>(n.s.) | n.s.<br>(n.s.) |
| | –10 | $p$<0.001<br>($p$<0.05) | n.s.<br>(n.s.) | | n.s.<br>(n.s.) | $p$<0.05<br>(n.s.) |
| | –8 | $p$<0.001<br>($p$<0.01) | n.s.<br>(n.s.) | n.s.<br>(n.s.) | | n.s.<br>(n.s.) |
| | vv6 | n.s.<br>($p$<0.001) | n.s.<br>(n.s.) | $p$<0.05<br>(n.s.) | n.s.<br>(n.s.) | |

**Time in bottom half (%) caused by 5CS+5DS**
**Kruskal-Wallis test: H (5, N= 56)=31.061, $p$ <0.001**
**Multiple Comparisons $p$ values (2-tailed) with Bonferroni correction**

| Fraction | 0 | –12 | –11 | –10 | –9 | SKE |
|---|---|---|---|---|---|---|
| | 8 | 10 | 10 | 10 | 10 | 8 |
| 0 | | $p$<0.001 | n.s. | $p$<0.001 | $p$<0.001 | n.s. |
| –12 | $p$<0.001 | | n.s. | n.s. | n.s. | n.s. |
| –11 | n.s. | n.s. | | n.s. | n.s. | n.s. |
| –10 | $p$<0.001 | n.s. | n.s. | | n.s. | $p$<0.05 |
| –9 | $p$<0.001 | n.s. | n.s. | n.s. | | n.s. |
| SKE | n.s. | n.s. | n.s. | $p$<0.05 | n.s. | |

**Freezing duration (%) caused by 5CS (5DS)**
**Tamhane test**
**Significance of homogeneity test of variance: *p* (5CS and 5DS)<0.05**

| | Fraction | 0 | –12 | –10 | –8 | –6 |
|---|---|---|---|---|---|---|
| N of biologically independent samples | | 6<br>(6) | 11<br>(9) | 9<br>(12) | 9<br>(10) | 7<br>(10) |
| | 0 | | $p<0.001$<br>($p<0.001$) | $p<0.001$<br>($p<0.001$) | $p<0.001$<br>($p<0.01$) | $p<0.01$<br>($p<0.001$) |
| | –12 | $p<0.001$<br>($p<0.001$) | | $p<0.05$<br>($p<0.01$) | $p<0.05$<br>(n.s.) | $p<0.05$<br>(n.s.) |
| | –10 | $p<0.001$<br>($p<0.001$) | $p<0.05$<br>($p<0.01$) | | n.s.<br>(n.s.) | n.s.<br>($p<0.001$) |
| | –8 | $p<0.001$<br>($p<0.01$) | $p<0.05$<br>(n.s.) | n.s.<br>(n.s.) | | n.s.<br>($p<0.01$) |
| | –6 | $p<0.01$<br>($p<0.001$) | $p<0.05$<br>(n.s.) | n.s.<br>($p<0.001$) | n.s.<br>($p<0.01$) | |

**Freezing duration (%) caused by 5CS+5DS**
**Tamhane test**
**Significance of homogeneity test of variance: p<0.05**

| Fraction | 0 | –12 | –11 | –10 | –9 | SKE |
|---|---|---|---|---|---|---|
| | 6 | 10 | 9 | 6 | 10 | 7 |
| 0 | | $p<0.01$ | $p<0.01$ | $p<0.001$ | $p<0.001$ | $p<0.001$ |
| –12 | $p<0.01$ | | n.s. | n.s. | n.s. | n.s. |
| –11 | $p<0.01$ | n.s. | | $p<0.001$ | $p<0.01$ | n.s. |
| –10 | $p<0.001$ | n.s. | $p<0.001$ | | n.s. | $p<0.01$ |
| –9 | $p<0.001$ | n.s. | $p<0.01$ | n.s. | | $p<0.01$ |
| SKE | $p<0.001$ | n.s. | n.s. | $p<0.01$ | $p<0.01$ | |

| | **Latency to upper half (s) caused by 5CS (5DS)<br>Tamhane test<br>Significance of homogeneity test of variance: *p* (5CS and 5DS)<0.05** | | | | | |
|---|---|---|---|---|---|---|
| | Fraction | 0 | –12 | –10 | –8 | –6 |
| N of biologically independent samples | | 6<br>(6) | 12<br>(11) | 11<br>(14) | 10<br>(11) | 8<br>(7) |
| | 0 | | $p$<0.001<br>($p$<0.001) | $p$<0.001<br>($p$<0.001) | $p$<0.001<br>($p$<0.001) | $p$<0.01<br>($p$<0.001) |
| | –12 | $p$<0.001<br>($p$<0.001) | | $p$<0.001<br>($p$<0.001) | $p$<0.05<br>(n.s.) | n.s.<br>(n.s.) |
| | –10 | $p$<0.001<br>($p$<0.001) | $p$<0.001<br>($p$<0.001) | | n.s.<br>(n.s.) | $p$<0.001<br>($p$<0.05) |
| | –8 | $p$<0.001<br>($p$<0.001) | $p$<0.05<br>(n.s.) | n.s.<br>(n.s.) | | n.s.<br>(n.s.) |
| | –6 | $p$<0.01<br>($p$<0.001) | n.s.<br>(n.s.) | $p$<0.001<br>($p$<0.05) | n.s.<br>(n.s.) | |

| **Latency to upper half (s) caused by 5CS+5DS<br>Tamhane test<br>Significance of homogeneity test of variance: *p*<0.05** | | | | | | |
|---|---|---|---|---|---|---|
| Fraction | 0 | –12 | –11 | –10 | –9 | SKE |
| | 6 | 10 | 10 | 6 | 10 | 6 |
| 0 | | $p$<0.001 | $p$<0.001 | $p$<0.001 | $p$<0.001 | $p$<0.001 |
| –12 | $p$<0.001 | | n.s. | $p$<0.05 | $p$<0.05 | $p$<0.01 |
| –11 | $p$<0.001 | n.s. | | $p$<0.001 | $p$<0.01 | $p$<0.01 |
| –10 | $p$<0.001 | $p$<0.05 | $p$<0.001 | | n.s. | $p$<0.001 |
| –9 | $p$<0.001 | $p$<0.05 | $p$<0.01 | n.s. | | $p$<0.001 |
| SKE | $p$<0.001 | $p$<0.01 | $p$<0.01 | $p$<0.001 | $p$<0.001 | |

**Supplementary Table 10:** Statistical analysis of alarm behaviors induced by different stimuli at the same concentration in figure 3a-d (sig. = significant, n.s. = not significant).

| **Erratic movements duration (%)** | | | | | | | | |
|---|---|---|---|---|---|---|---|---|
| Concentration (lg[M]) | –12 | | –10 | | –8 | | –6 | |
| Compound | 5CS | 5DS | 5CS | 5DS | 5CS | 5DS | 5CS | 5DS |
| N of biologically independent samples | 8 | 11 | 9 | 12 | 10 | 9 | 16 | 14 |
| Tests of normality (sig. of Shapiro-Wilk) | 0.268 | 0.551 | 0.901 | 0.273 | 0.472 | 0.243 | 0.587 | 0.062 |
| Statistical analyses p values (2-tailed) | Independent samples t-test | | | | | | | |
| | n.s. | | *p*<0.01 | | *p*<0.001 | | n.s. | |
| **Time in bottom half (%)** | | | | | | | | |
| Concentration (lg[M]) | –12 | | –10 | | –8 | | –6 | |
| Compound | 5CS | 5DS | 5CS | 5DS | 5CS | 5DS | 5CS | 5DS |
| N of biologically independent samples | 20 | 20 | 20 | 20 | 20 | 20 | 20 | 20 |
| Tests of normality (sig. of Shapiro-Wilk) | 0.029 | 0.007 | 0.019 | 0.005 | 0.051 | 0.000 | 0.17 | 0.000 |
| Statistical analyses p values (2-tailed) | Mann-Whitney U test | | | | | | | |
| | n.s. | | n.s. | | n.s. | | *p*<0.001 | |
| **Freezing duration (%)** | | | | | | | | |
| Concentration (lg[M]) | –12 | | –10 | | –8 | | –6 | |
| Compound | 5CS | 5DS | 5CS | 5DS | 5CS | 5DS | 5CS | 5DS |
| N of biologically independent samples | 11 | 9 | 9 | 12 | 9 | 10 | 7 | 10 |
| Tests of normality (sig. of Shapiro-Wilk) | 0.810 | 0.607 | 0.619 | 0.496 | 0.716 | 0.360 | 0.801 | 0.203 |
| Statistical analyses p values (2-tailed) | Independent samples t-test | | | | | | | |
| | *p*<0.001 | | n.s. | | *p*<0.05 | | *p*<0.001 | |

| Latency to upper half (s) | | | | | | | | |
|---|---|---|---|---|---|---|---|---|
| Concentration (lg[M]) | –12 | | –10 | | –8 | | –6 | |
| Compound | 5CS | 5DS | 5CS | 5DS | 5CS | 5DS | 5CS | 5DS |
| N of biologically independent samples | 12 | 11 | 11 | 14 | 10 | 11 | 8 | 7 |
| Tests of normality (sig. of Shapiro-Wilk) | 0.518 | 0.958 | 0.374 | 0.117 | 0.086 | 0.294 | 0.505 | 0.570 |
| Statistical analyses p values (2-tailed) | Independent samples t-test | | | | | | | |
| | $p$<0.001 | | $p$<0.001 | | n.s. | | n.s. | |

**Supplementary Table 11:** Statistical analysis of whole-body cortisol level in figure 4 (n.s. = not significant).

**Whole-body cortisol level exposed to 5CS (5DS)**
**Kruskal-Wallis test of 5CS: H (4, N= 68)=52.360, *p* <0.001; Kruskal-Wallis test of 5DS: H (4, N= 67)=35.084, *p* <0.001**
**Multiple Comparisons *p* values (2-tailed) with Bonferroni correction**

| | Fraction | 0 | –12 | –10 | –8 | –6 |
|---|---|---|---|---|---|---|
| N of biologically independent samples | | 15 (15) | 12 (12) | 12 (13) | 15 (13) | 14 (14) |
| | 0 | | n.s. (n.s.) | $p$<0.01 (n.s.) | $p$<0.001 ($p$<0.01) | $p$<0.001 ($p$<0.01) |
| | –12 | n.s. (n.s.) | | n.s. (n.s.) | n.s. ($p$<0.001) | $p$<0.001 ($p$<0.001) |
| | –10 | $p$<0.01 (n.s.) | n.s. (n.s.) | | n.s. (n.s.) | $p$<0.05 (n.s.) |
| | –8 | $p$<0.001 (p<0.01) | n.s. ($p$<0.001) | n.s. (n.s.) | | $p$<0.05 (n.s.) |
| | –6 | $p$<0.001 (p<0.01) | $p$<0.001 ($p$<0.001) | $p$<0.05 (n.s.) | $p$<0.05 (n.s.) | |

**Supplementary Table 12:** Statistical analysis of whole-body cortisol level induced by different stimuli at the same concentration in figure 4 (sig. = significant, n.s. = not significant).

| Whole-body cortisol level | | | | | | | | | | |
|---|---|---|---|---|---|---|---|---|---|---|
| Concentration (lg[M]) | 0 | | –12 | | –10 | | –8 | | –6 | |
| Compound | 5CS | 5DS | 5CS | 5DS | 5CS | 5DS | 5CS | 5DS | 5CS | 5DS |
| N of biologically independent samples | 15 | 15 | 12 | 12 | 12 | 13 | 15 | 13 | 14 | 14 |
| Tests of normality (sig. of Shapiro-Wilk) | 0.891 | 0.891 | 0.053 | 0.052 | 0.761 | 0.416 | 0.005 | 0.027 | 0.257 | 0.197 |
| Statistical analyses *p* values (2-tailed) | Independent samples t-test | | | | | | Mann-Whitney U test | | Independent samples t-test | |
| | n.s. | | $p$<0.001 | | $p$<0.001 | | n.s. | | $p$<0.001 | |

**Supplementary Table 13:** Statistical analysis of zebrafish anti-predation behavior phenotypes triggered by SKE dilutions (sig. = significant, n.s. = not significant, Con. = Control, Exp. = Experiment).

| **Erratic movements duration (%)** | | | | | | | | | | | | |
|---|---|---|---|---|---|---|---|---|---|---|---|---|
| lg[SKE dilution] | 6 | | 5 | | 4 | | 3 | | 2 | | 1 | |
| Group | Con. | Exp. | Con. | Exp. | Con. | Exp. | Con. | Exp. | Con. | Exp. | Con. | Exp. |
| N of biologically independent samples | 15 | 15 | 15 | 15 | 15 | 15 | 15 | 15 | 15 | 15 | 15 | 15 |
| Tests of normality (sig. of Shapiro-Wilk) | 0.001 | 0.215 | 0.001 | 0.013 | 0.248 | 0.316 | 0.010 | 0.015 | 0.775 | 0.022 | 0.235 | 0.008 |
| Statistical analyses *p* values (2-tailed) | Mann-Whitney U test | | | | Independent samples t-test | | Mann-Whitney U test | | | | | |
| | n.s. | | n.s. | | n.s. | | $p$<0.001 | | $p$<0.001 | | n.s. | |
| **Time in bottom half (%)** | | | | | | | | | | | | |
| lg[SKE dilution] | 6 | | 5 | | 4 | | 3 | | 2 | | 1 | |
| Group | Con. | Exp. | Con. | Exp. | Con. | Exp. | Con. | Exp. | Con. | Exp. | Con. | Exp. |
| N of biologically independent samples | 12 | 12 | 12 | 12 | 12 | 12 | 12 | 12 | 12 | 12 | 12 | 12 |
| Tests of normality (sig. of Shapiro-Wilk) | 0.204 | 0.366 | 0.085 | 0.293 | 0.806 | 0.267 | 0.046 | 0.816 | 0.107 | 0.294 | 0.846 | 0.552 |
| Statistical analyses *p* values (2-tailed) | Independent samples t-test | | | | | | Mann-Whitney U test | | Independent samples t-test | | | |
| | n.s. | | n.s. | | n.s. | | $p$<0.001 | | n.s. | | n.s. | |
| **Freezing duration (%)** | | | | | | | | | | | | |
| lg[SKE dilution] | 6 | | 5 | | 4 | | 3 | | 2 | | 1 | |
| Group | Con. | Exp. | Con. | Exp. | Con. | Exp. | Con. | Exp. | Con. | Exp. | Con. | Exp. |
| N of biologically independent samples | 9 | 9 | 9 | 9 | 9 | 9 | 9 | 9 | 9 | 9 | 9 | 9 |
| Tests of normality (sig. of Shapiro-Wilk) | 0.767 | 0.896 | – | 0.223 | 0.407 | 0.103 | 0.533 | 0.103 | – | 0.379 | 0.126 | 0.278 |

| | | | | | | | | | | | | |
|---|---|---|---|---|---|---|---|---|---|---|---|---|
| Statistical analyses p values (2-tailed) | Independent samples t-test<br>n.s. | | n.s. | | n.s. | | Mann-Whitney U test<br>p<0.001 | | Independent samples t-test<br>n.s. | | n.s. | |
| **Latency to upper half (s)** | | | | | | | | | | | | |
| lg[SKE dilution] | 6 | | 5 | | 4 | | 3 | | 2 | | 1 | |
| Group | Con. | Exp. | Con. | Exp. | Con. | Exp. | Con. | Exp. | Con. | Exp. | Con. | Exp. |
| N of biologically independent samples | 7 | 7 | 7 | 7 | 7 | 7 | 7 | 7 | 7 | 7 | 7 | 7 |
| Tests of normality (sig. of Shapiro-Wilk) | 0.888 | 0.605 | 0.388 | 0.162 | 0.451 | 0.867 | 0.319 | 0.872 | 0.341 | 0.207 | 0.171 | 0.572 |
| Statistical analyses *p* values (2-tailed) | Independent samples t-test<br>n.s. | | n.s. | | n.s. | | *p*<0.001 | | *p*<0.001 | | *p*<0.001 | |
| **Number of entyies to the bottom** | | | | | | | | | | | | |
| lg[SKE dilution] | 6 | | 5 | | 4 | | 3 | | 2 | | 1 | |
| Group | Con. | Exp. | Con. | Exp. | Con. | Exp. | Con. | Exp. | Con. | Exp. | Con. | Exp. |
| N of biologically independent samples | 15 | 15 | 15 | 15 | 15 | 15 | 15 | 15 | 15 | 15 | 15 | 15 |
| Tests of normality (sig. of Shapiro-Wilk) | 0.531 | 0.038 | 0.398 | 0.649 | 0.311 | 0.557 | 0.511 | 0.289 | 0.006 | 0.001 | 0.055 | 0.867 |
| Statistical analyses p values (2-tailed) | Mann-Whitney U test<br>n.s. | | Independent samples t-test<br>n.s. | | n.s. | | n.s. | | Mann-Whitney U test<br>n.s. | | Independent samples t-test<br>n.s. | |
| **Distance traveled in bottom (m)** | | | | | | | | | | | | |
| lg[SKE dilution] | 6 | | 5 | | 4 | | 3 | | 2 | | 1 | |
| Group | Con. | Exp. | Con. | Exp. | Con. | Exp. | Con. | Exp. | Con. | Exp. | Con. | Exp. |
| N of biologically independent samples | 15 | 15 | 15 | 15 | 15 | 15 | 15 | 15 | 15 | 15 | 15 | 15 |
| Tests of normality (sig. of Shapiro-Wilk) | 0.076 | 0.061 | 0.055 | 0.310 | 0.187 | 0.069 | 0.061 | 0.205 | 0.019 | 0.289 | 0.084 | 0.197 |
| Statistical analyses *p* values (2-tailed) | Independent samples t-test<br>n.s. | | n.s. | | n.s. | | Mann-Whitney U test<br>n.s. | | Independent samples t-test<br>n.s. | | n.s. | |

| | | | | | | | | | | | | |
|---|---|---|---|---|---|---|---|---|---|---|---|---|
| **Average velocity (cm/s)** | | | | | | | | | | | | |
| lg[SKE dilution] | 6 | | 5 | | 4 | | 3 | | 2 | | 1 | |
| Group | Con. | Exp. | Con. | Exp. | Con. | Exp. | Con. | Exp. | Con. | Exp. | Con. | Exp. |
| N of biologically independent samples | 15 | 15 | 15 | 15 | 15 | 15 | 15 | 15 | 15 | 15 | 15 | 15 |
| Tests of normality (sig. of Shapiro-Wilk) | 0.298 | 0.134 | 0.376 | 0.857 | 0.339 | 0.126 | 0.029 | 0.494 | 0.522 | 0.588 | 0.318 | 0.025 |
| Statistical analyses *p* values (2-tailed) | Independent samples t-test | | | | | | Mann-Whitney U test | | Independent samples t-test | | Mann-Whitney U test | |
| | n.s. | | n.s. | | n.s. | | n.s. | | n.s. | | n.s. | |
| **Total distance travelled (m)** | | | | | | | | | | | | |
| lg[SKE dilution] | 6 | | 5 | | 4 | | 3 | | 2 | | 1 | |
| Group | Con. | Exp. | Con. | Exp. | Con. | Exp. | Con. | Exp. | Con. | Exp. | Con. | Exp. |
| N of biologically independent samples | 15 | 15 | 15 | 15 | 15 | 15 | 15 | 15 | 15 | 15 | 15 | 15 |
| Tests of normality (sig. of Shapiro-Wilk) | 0.080 | 0.081 | 0.164 | 0.047 | 0.061 | 0.069 | 0.150 | 0.850 | 0.002 | 0.002 | 0.054 | 0.010 |
| Statistical analyses *p* values (2-tailed) | Independent samples t-test | | Mann-Whitney U test | | Independent samples t-test | | | | Mann-Whitney U test | | | |
| | n.s. | | n.s. | | n.s. | | n.s. | | n.s. | | n.s. | |
| **Time spent (bottom/top)** | | | | | | | | | | | | |
| lg[SKE dilution] | 6 | | 5 | | 4 | | 3 | | 2 | | 1 | |
| Group | Con. | Exp. | Con. | Exp. | Con. | Exp. | Con. | Exp. | Con. | Exp. | Con. | Exp. |
| N of biologically independent samples | 12 | 12 | 12 | 12 | 12 | 12 | 12 | 12 | 12 | 12 | 12 | 12 |
| Tests of normality (sig. of Shapiro-Wilk) | 0.013 | 0.973 | 0.307 | 0.324 | 0.011 | 0.391 | 0.417 | 0.518 | 0.021 | 0.004 | 0.294 | 0.224 |
| Statistical analyses *p* values (2-tailed) | Mann-Whitney U test | | Independent samples t-test | | Mann-Whitney U test | | Independent samples t-test | | Mann-Whitney U test | | Independent samples t-test | |
| | $p<0.01$ | | $p<0.01$ | | n.s. | | $p<0.001$ | | n.s. | | $p<0.001$ | |
| **Entries (bottom/top)** | | | | | | | | | | | | |

| lg[SKE dilution] | 6 | | 5 | | 4 | | 3 | | 2 | | 1 | |
|---|---|---|---|---|---|---|---|---|---|---|---|---|
| Group | Con. | Exp. | Con. | Exp. | Con. | Exp. | Con. | Exp. | Con. | Exp. | Con. | Exp. |
| N of biologically independent samples | 12 | 12 | 12 | 12 | 12 | 12 | 12 | 12 | 12 | 12 | 12 | 12 |
| Tests of normality (sig. of Shapiro-Wilk) | 0.546 | 0.227 | 0.000 | 0.052 | 0.324 | 0.080 | 0.014 | 0.061 | 0.003 | 0.232 | 0.370 | 0.006 |
| Statistical analyses *p* values (2-tailed) | Independent samples t-test | | Mann-Whitney U test | | Independent samples t-test | | Mann-Whitney U test | | | | | |
| | n.s. | | n.s. | | n.s. | | n.s. | | n.s. | | n.s. | |
| **Average bottom entry duration (s)** | | | | | | | | | | | | |
| lg[SKE dilution] | 6 | | 5 | | 4 | | 3 | | 2 | | 1 | |
| Group | Con. | Exp. | Con. | Exp. | Con. | Exp. | Con. | Exp. | Con. | Exp. | Con. | Exp. |
| N of biologically independent samples | 10 | 10 | 10 | 10 | 10 | 10 | 10 | 10 | 10 | 10 | 10 | 10 |
| Tests of normality (sig. of Shapiro-Wilk) | 0.272 | 0.698 | 0.002 | 0.000 | 0.037 | 0.622 | 0.030 | 0.633 | 0.355 | 0.035 | 0.319 | 0.011 |
| Statistical analyses *p* values (2-tailed) | Independent samples t-test | | Mann-Whitney U test | | | | | | | | | |
| | $p$<0.05 | | n.s. | | n.s. | | $p$<0.001 | | n.s. | | $p$<0.05 | |
| **Distance traveled (bottom/top)** | | | | | | | | | | | | |
| lg[SKE dilution] | 6 | | 5 | | 4 | | 3 | | 2 | | 1 | |
| Group | Con. | Exp. | Con. | Exp. | Con. | Exp. | Con. | Exp. | Con. | Exp. | Con. | Exp. |
| N of biologically independent samples | 12 | 12 | 12 | 12 | 12 | 12 | 12 | 12 | 12 | 12 | 12 | 12 |
| Tests of normality (sig. of Shapiro-Wilk) | 0.899 | 0.627 | 0.471 | 0.073 | 0.552 | 0.271 | 0.443 | 0.316 | 0.051 | 0.216 | 0.272 | 0.350 |
| Statistical analyses *p* values (2-tailed) | Independent samples t-test | | | | | | | | | | | |
| | $p$<0.001 | | $p$<0.05 | | n.s. | | $p$<0.01 | | n.s. | | $p$<0.001 | |
